# A large-scale foundation model for bulk transcriptomes

**DOI:** 10.1101/2025.06.11.659222

**Authors:** Boming Kang, Rui Fan, Meizheng Yi, Chunmei Cui, Qinghua Cui

**Author notes:** These authors contributed equally to this work (Boming Kang, Rui Fan). To whom the correspondence should be addressed: Qinghua Cui.

## Abstract

Large language models (LLMs) have emerged as powerful foundation models leading to breakthroughs in transcriptome analysis. However, current RNA-seq foundation models are exclusively pretrained on sparse single-cell RNA-seq (scRNA-seq) data, which typically detects only ∼3000 genes per cell. This thus creates a critical gap in models specifically designed for bulk transcriptomes, a fundamentally different modality capable of profiling ∼16,000 genes per sample. Here we propose BulkFormer, a large-scale foundation model for bulk transcriptome analysis. With 150 million parameters covering about 20,000 protein-coding genes, BulkFormer is pretrained on over 500,000 human bulk transcriptomic profiles. BulkFormer incorporates a hybrid encoder architecture, combining a graph neural network to capture explicit gene-gene interactions and a performer module to model global expression dependencies. As a result, despite incurring much lower training costs than scRNA-seq foundation models, BulkFormer consistently outperforms them in all six downstream tasks: transcriptome imputation, disease annotation, prognosis modeling, drug response prediction, compound perturbation simulation, and gene essentiality scoring. Notably, BulkFormer not only enhances the discovery of novel clinical biomarkers but also uncovers latent disease mechanisms by imputing biologically meaningful gene expression. Collectively, these results demonstrate BulkFormer’s power as a versatile and robust framework for bulk transcriptome modeling and biomedical discovery, bridging a critical gap in the current foundation model landscape.

## Introduction

Large-scale pretrained language models represent a revolutionary breakthrough in the field of natural language processing (NLP) in recent years^1^. Similar to natural language, DNA, RNA, and protein sequences in life sciences can also be regarded as biological languages, leading to the development of a series of large-scale pretrained biological language models^2^, such as DNA-BERT^3^, RNA-FM^4^, and ESM2^5^. Unlike biological sequences, gene expression profiles derived from transcriptomics encode the biological information of life systems, serving as a functional language that reflects the physiological state of the living organisms^6^. For example, patient prognosis can be predicted using the expression patterns of disease-associated biomarkers^7^. Transcriptomic sequencing technologies can be broadly classified into bulk RNA sequencing and single-cell RNA sequencing (scRNA-seq). Bulk RNA-seq measures the average gene expression across a population of cells, thus providing a global but low-resolution view of transcriptional activity. In contrast, scRNA-seq captures gene expression at single-cell resolution, enabling the identification of cellular heterogeneity and rare cell types. Therefore, the substantially larger scale of gene expression data generated by scRNA-seq compared to bulk RNA-seq has driven the development of a series of foundation models specifically pretrained on single-cell transcriptomic data, including Geneformer^8^, scGPT^9^, scFoundation^10^, GeneCompass^11^, and scLong^12^. Single-cell large language models (scLLMs) have demonstrated the ability to extract high-quality cellular and gene-level transcriptomic representations, enabling state-of-the-art performance in diverse downstream single-cell tasks such as cell annotation, drug response prediction, perturbation effect prediction, and gene module inference^13,14^. Although scRNA-seq provides single-cell resolution, its inherent sparsity, defined as the limited detection of gene expression per cell^15^, presents difficulties for downstream tasks requiring comprehensive transcriptomic coverage, such as disease subtype classification and prognostic modeling^16^. In contrast, bulk RNA-seq offers more comprehensive and stable gene expression measurements across samples, making it well-suited for system- or tissue-level analyses. However, large-scale models pretrained on bulk transcriptomic data still remain unavailable, highlighting a critical gap in the fields of transcriptomic modeling.

Given the characteristics of scRNA-seq data, existing scLLM models have adopted various training strategies. Geneformer ranks normalized gene expression values within each cell and is trained to predict the rank value of masked genes. scGPT discretize gene expression into bins and predict the bin membership of masked genes. While these approaches help mitigate data noise, they reduce the resolution of expression modeling and may impair performance in downstream tasks. To address this limitation, scFoundation and scLong directly predict the continuous expression values of masked genes, thereby improving modeling resolution. Distinctively, GeneCompass employs a multitask pretraining strategy with a dual-decoder architecture: one decoder predicts the gene identity at masked positions, and the other predicts the corresponding expression values. However, the encoder inputs of these models typically include only the top expressed few thousand genes in each sample, while the remaining large-number of genes, which are failed to be detected due to technical limitations, are assigned zero expression values. Although this strategy is appropriate for handling the sparsity of scRNA-seq data, it prevents the model from learning the complete set of gene-gene relationships across the whole transcriptome. As a result, these scLLMs are not well suited for bulk RNA-seq data and its associated downstream tasks.

In this study, we focused on bulk RNA-seq modeling and proposed BulkFomer, a large-scale foundation model with approximately 150 million parameters, covering around 20,000 protein-coding genes. To enable large-scale pretraining, we curated and standardized approximately 520,000 bulk RNA-seq gene expression profiles from public databases. To more effectively model bulk RNA-seq data, we developed a hybrid encoder architecture that integrates both graph neural networks (GNN) for capturing explicit gene-gene relationships from a biological knowledge graph while employing attention mechanisms to learn implicit transcriptional dependencies across the entire transcriptome.

To verify BulkFormer’s power, we performed extensive benchmarking across six critical downstream tasks, including transcriptome imputation, disease annotation, prognosis modeling, drug response prediction, compound perturbation prediction, and gene essentiality prediction. As a result, BulkFormer outperforms existing scLLMs in all tasks. Notably, when applied to clinical samples, BulkFormer successfully reconstructed missing gene expression values, enabling the discovery of a series of previously unrecognized prognostic biomarkers. These results collectively establish BulkFormer as a powerful and versatile tool for bulk RNA-seq modeling and analysis. This work not only advances the development of foundation models for bulk transcriptomics but also opens new avenues for their biomedical applications.

## Results

### Overview of BulkFormer

To construct a large-scale dataset for BulkFormer pretraining, we first curated PreBULK, a comprehensive bulk transcriptomic dataset assembled from public repositories including Gene Expression Omnibus (GEO)^17^ and ARCHS4^18^, comprising 522,769 gene expression profiles for 20,010 protein-coding genes. BulkFormer was pretrained using a masked language modeling (MLM) strategy inspired by BERT^19^, in which 15% of gene expression values in the input transcriptome are randomly masked. The model is then trained to reconstruct the masked values by minimizing the mean squared error (MSE) loss between the predicted and true expression values, thereby updating model parameters (**Fig. 1a**).

**Fig. 1.**
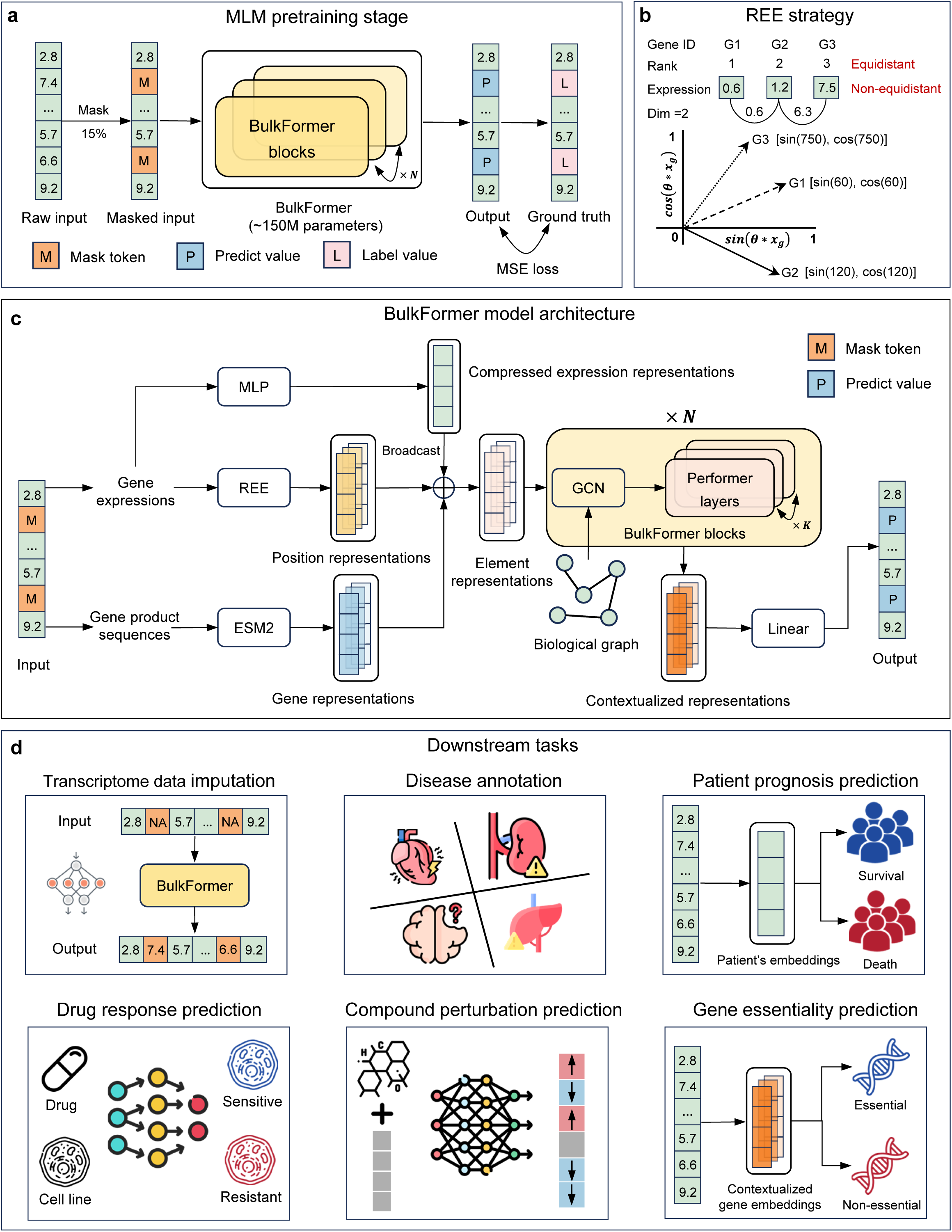
Overview of the BulkFormer framework and its applications. **(a)** The pretraining phase of BulkFormer adopts a masked language modeling (MLM) strategy, in which approximately 15% of gene expression values in each input sample are randomly masked. The model is trained to predict the masked values based on context, and parameters are optimized using the mean squared error (MSE) loss between the predicted and the true values. **(b)** Schematic illustration of the rotary expression embedding (REE) strategy for encoding gene expression values. **(c)** Model architecture of BulkFormer. ESM2 was used to extract sequence-based embeddings of canonical protein products, serving as initial representations for individual gene tokens. Each gene’s expression value was treated as a positional token and encoded using rotary position embedding to capture continuous expression information. Simultaneously, a MLP module compressed the global expression vector into a sample-level embedding. These three representations were fused via element-wise summation to form the final model input. The core of BulkFormer consists of stacked blocks, each containing a graph convolutional network layer to model gene–gene relationships followed by *K* Performer layers to capture long-range interactions. After *N* such blocks, contextualized gene embeddings are output and passed through a linear projection layer to predict gene expression levels. **(d)** Downstream applications of BulkFormer include transcriptome imputation, disease annotation, prognosis modeling, drug response prediction, compound perturbation prediction, and gene essentiality prediction.

To accommodate the specific modality of bulk transcriptomic data, we designed a distinct model architecture different from existing scLLMs. For each input sample, we utilized ESM2, a large-scale pretrained protein language model, to extract sequence-derived embeddings of the canonical protein products. These embeddings were used as the initial representations for individual gene tokens, based on the rationale that proteins, as the primary functional products of genes, play central roles in biological processes and directly reflect gene function at the molecular level^20^. The rank of gene expression values can be regarded as a form of gene ordering within each sample, analogous to positional encoding in LLMs. However, rank-based values are evenly spaced and fail to capture the relative magnitudes between gene expression levels. Inspired by rotary position encoding (ROPE)^21^, we propose a rotary expression embedding (REE) strategy to encode gene expression values as positional representations, preserving both their magnitude and continuity (**See Methods for details**). The embeddings generated by REE retain the relative relationships between gene expression levels without requiring additional training, offering strong stability and interpretability (**Fig. 1b**). In parallel, a multilayer perceptron (MLP) module was employed to compress the input expression vector into a global sample-level embedding. These three representations were then integrated via element-wise summation to form the final input representations. The core architecture of BulkFormer consists of stacked BulkFormer blocks, each composed of one graph convolutional network (GCN)^22^ layer followed by *K* Performer^23^ layers. The GCN module leverages a prior biological knowledge graph to capture explicit gene-gene relationships, while the performer, a scalable variant of the transformer^24^ is employed to model implicit gene interactions. The Performer approximates self-attention with linear complexity, making it particularly well-suited for processing high-dimensional bulk transcriptomic inputs. After passing through *N* BulkFormer blocks, the model yields contextualized gene embeddings, which are projected through a linear layer to generate scalar gene expression predictions **(Fig. 1c)**. Detailed hyperparameter settings for BulkFormer, as well as results from ablation studies, were provided in the supplementary information. To assess BulkFormer’s ability, we evaluated its performance across a series of canonical bulk transcriptome tasks, including transcriptome imputation, disease annotation, prognosis modeling, drug response prediction, compound perturbation prediction, and gene essentiality prediction **(Fig. 1d)**. In all tasks, BulkFormer consistently outperformed existing baseline models, highlighting its capacity and versatility for bulk transcriptome modeling (**Supplementary Table 1**). Notably, pretraining BulkFormer on bulk transcriptomic data is an efficient and computationally economical approach. BulkFormer requires only 1% to 10% of the training time (measured by the average single-GPU epoch duration) compared to existing scLLMs, while still effectively capture gene–gene relationships and generate high-quality gene-level and transcriptome-level representations for downstream tasks (**Supplementary Table 2**).

### Pretraining on large-scale bulk transcriptomes enables biologically meaningful embeddings

Owing to technical limitations such as low mRNA capture efficiency and sequencing depth per cell, scRNA-seq typically detects only 500 to 5,000 genes per cell. By comparison, bulk RNA-seq routinely quantifies the expression of over 15,000 to 20,000 protein-coding genes per sample, offering a more complete view of the transcriptome. In bulk RNA-seq, the expression level of each gene represents an aggregate signal, reflecting the average expression across all constituent cells within the sampled tissue (**Fig. 2a**). We further compared the sparsity of PreBULK with that of the single-cell dataset sourced from Tabula Sapiens^25^ by evaluating the number of non-zero protein-coding genes detected per gene expression profile. On average, PreBULK captured expression for 16,606 protein-coding genes per sample, whereas the single-cell dataset detected only 3,050 genes per cell (**Fig. 2b**). This stark contrast underscores the difference in data sparsity between the two modalities, highlighting the substantially more complete transcriptomic coverage provided by bulk RNA-seq. PreBULK encompasses bulk transcriptomic profiles from nine major human physiological systems and their associated tissues, enabling BulkFormer to learn comprehensive representations of the human bulk transcriptome (**Fig. 2c**). During the pretraining stage, BulkFormer was trained for a total of 29 epochs, with the MSE loss on the test set steadily decreasing from an initial value of 7.5582 to 0.2427, indicating convergence and completion of model training (**Fig. 2d**). To visualize the learned representations, we extracted the gene token embeddings from BulkFormer’s embedding layer and applied t-distributed stochastic neighbor embedding (t-SNE)^26^ for dimensionality reduction. The resulting two-dimensional map revealed that genes with distinct average expression levels were grouped into different clusters **(Fig. 2e)**. To evaluate the biological relevance of the learned embeddings, we performed K-means clustering on all gene embeddings, partitioning them into 10 clusters **(Fig. 2f)**. Gene Ontology (GO)^27^ enrichment analysis of each cluster revealed a varying number of enriched terms, ranging from 69 to 1,388 across clusters **(Fig. 2g)**, with distinct biological functions observed in different gene groups **(Supplementary Fig. 1)**. For example, cluster 6 genes were predominantly associated with DNA replication and cell cycle, whereas cluster 9 genes were enriched in immune-related functions **(Fig. 2h)**. Consistently, Kyoto Encyclopedia of Genes and Genomes (KEGG)^28^ pathway enrichment analysis showed that the number of enriched pathways per cluster ranged from 12 to 172 **(Fig. 2i)**, and the functional annotations were likewise cluster-specific **(Supplementary Fig. 2)**. For instance, cluster 2 genes were enriched in innate immune signaling pathways, while cluster 7 genes were enriched in cancer-related pathways **(Fig. 2j)**.

**Fig. 2.**
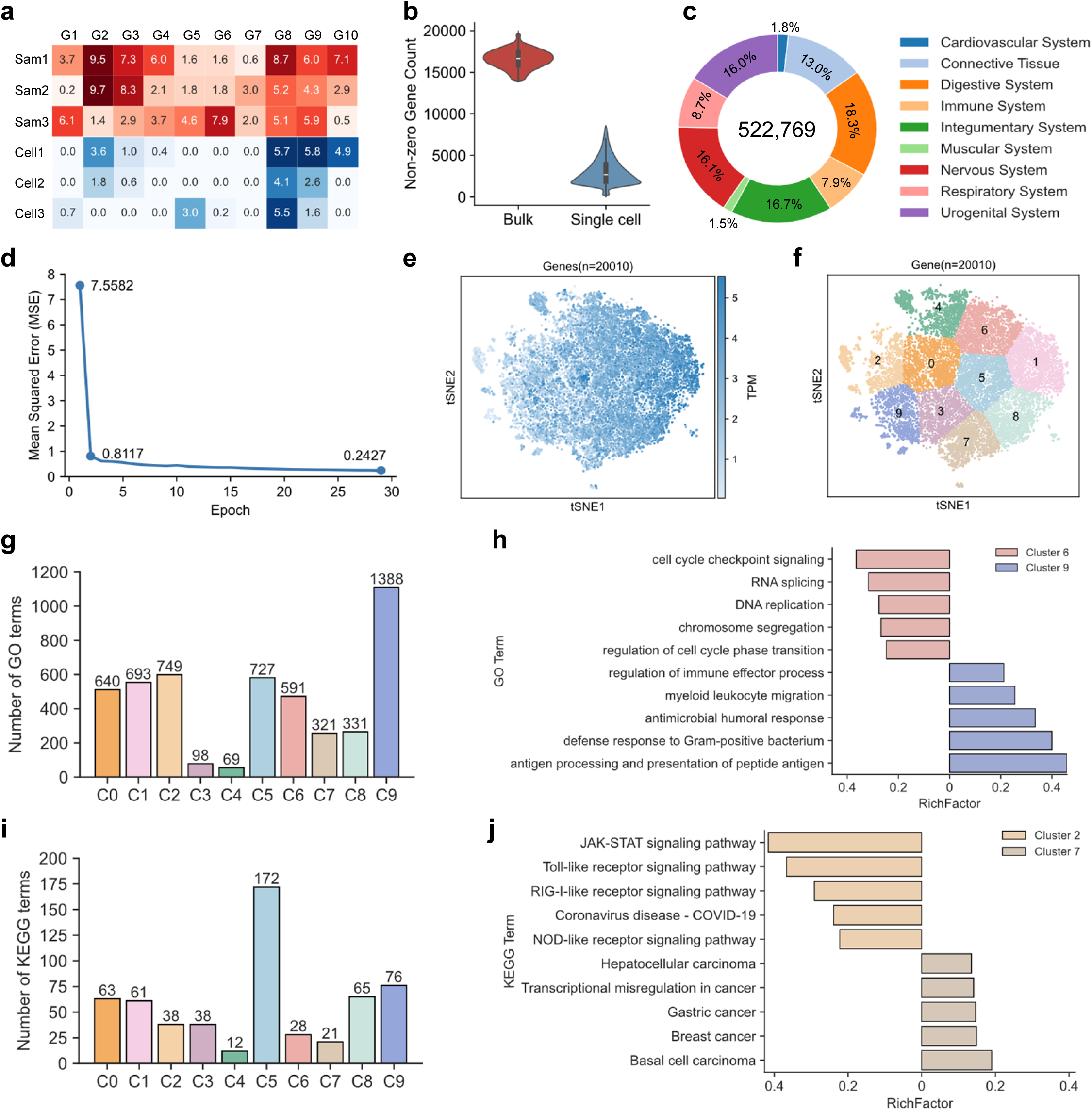
BulkFormer pretrained on large-scale bulk transcriptomes enables biologically meaningful embeddings. **(a)** Schematic illustration comparing the modality characteristics of single-cell RNA-seq (scRNA-seq) and bulk RNA-seq. **(b)** Comparison of the number of detected genes per sample between BulkFormer’s pretraining bulk RNA-seq dataset and the single-cell transcriptomes from the Tabula Sapiens database. **(c)** Distribution of human organ systems represented in the bulk transcriptomic dataset used for BulkFormer pretraining. **(d)** Loss curve on the held-out test set during BulkFormer pretraining. **(e)** t-SNE visualization of gene embeddings extracted from BulkFormer’s embedding layer. Color intensity reflects the average expression level (TPM: transcripts per million) of each gene across the training set. **(f)** K-means clustering of gene embeddings (k = 10), followed by t-SNE dimensionality reduction for visualization. **(g)** Number of significantly enriched Gene Ontology (GO) terms associated with genes in each cluster. **(h)** Comparative GO term enrichment results for Gene Cluster 6 and Gene Cluster 9. **(i)** Number of significantly enriched KEGG pathways associated with genes in each cluster. **(j)** Comparative KEGG term enrichment results for Gene Cluster 2 and Gene Cluster 7.

Together, these results demonstrate that BulkFormer not only successfully converges during pretraining but also produces biologically meaningful gene embeddings that reflect functional diversity within the human transcriptome.

### BulkFormer enables context-aware imputation of missing values from bulk transcriptomes

BulkFormer was pretrained using a MLM strategy, in which a subset of gene expression values is masked and subsequently reconstructed based on contextual information from the remaining genes. This naturally lends BulkFormer to the task of imputing missing values in bulk transcriptomic datasets. To evaluate this capability, we randomly masked 15% of the gene expression values in each sample within an independent test set to simulate realistic dropout events, and compared the imputation performance of BulkFormer against a range of scLLMs. Geneformer, GeneCompass, and scGPT were not directly pretrained to model gene expression values, and therefore cannot perform transcriptome imputation tasks. We therefore included the variational autoencoder (VAE)^29^ as an additional baseline model for comparison. As a result, BulkFormer achieved the highest imputation accuracy, yielding a Pearson correlation coefficient (PCC) of r = 0.954 between imputed and true values **(Fig. 3a)**. The next-best model was the VAE (r = 0.806), followed by simple imputation methods using the gene-wise mean (r = 0.754) or median (r = 0.743) across the training set. Models pretrained on scRNA-seq data, including scFoundation and scLong, performed substantially worse (r < 0.150), likely due to their incompatibility with the bulk data modality.

**Fig. 3.**
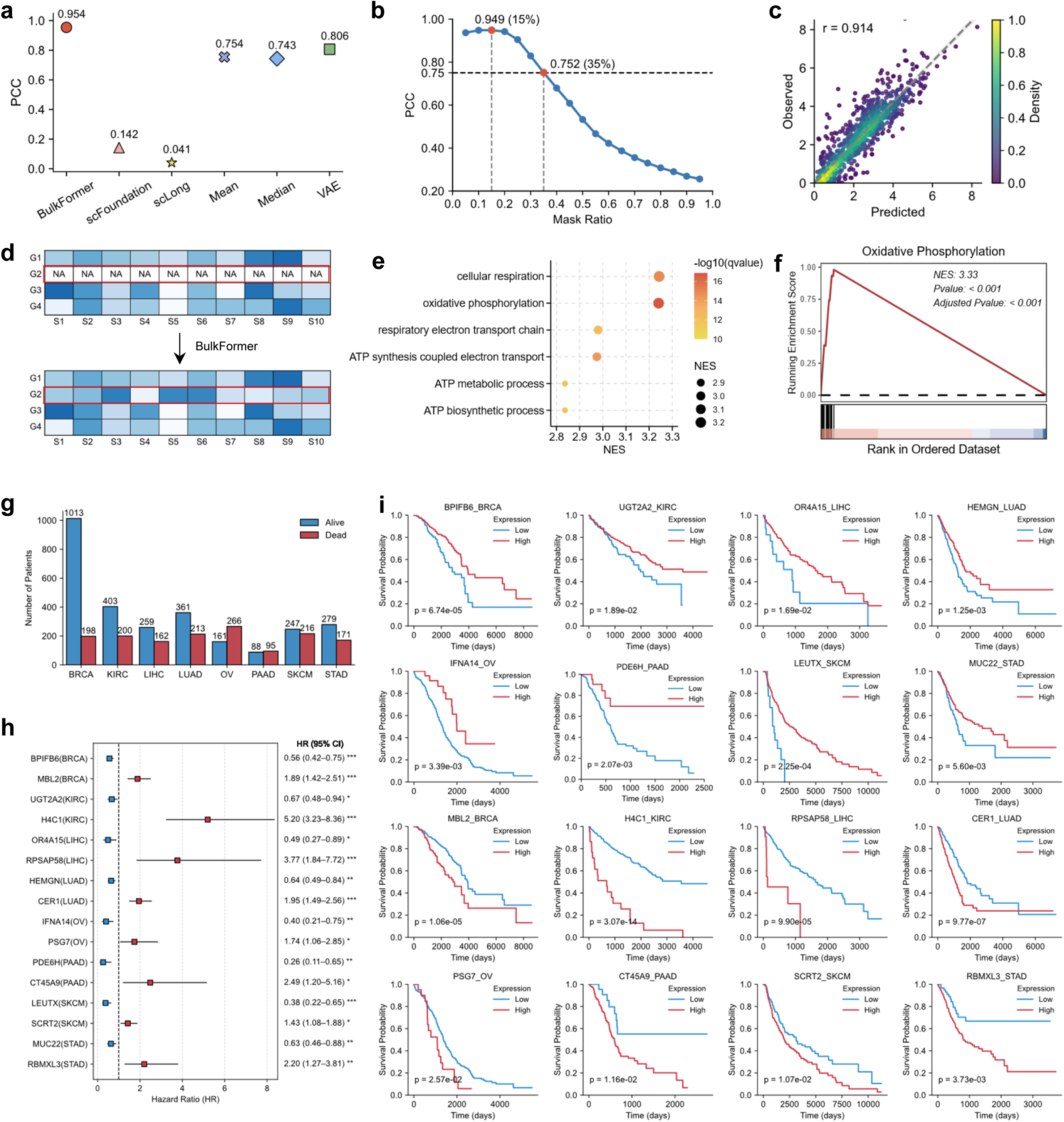
BulkFormer enables context-aware imputation of missing values from bulk transcriptomes. **(a)** Performance comparison between BulkFormer and baseline models on the transcriptome imputation task. **(b)** Effect of varying gene expression missing rates on the imputation performance of BulkFormer. **(c)** The imputation results of BulkFormer on transcriptomic data from the TCGA database. **(d)** Schematic illustration of BulkFormer’s ability to contextually impute gene expression values even for genes that are completely missing from a sample. **(e–f)** GSEA results of differentially expressed genes newly identified after BulkFormer-based imputation. **(g)** Distribution of survival and death outcomes in eight selected cancer patient cohorts from the TCGA database. **(h)** Prognostic biomarkers newly discovered via BulkFormer-based imputation, including both risk and protective factors. Genes with HR > 1 are defined as risk factors, and those with HR < 1 are defined as protective factors. **(i)** Kaplan-Meier survival curves for cancer patients stratified by expression levels of newly discovered prognostic biomarkers imputed by BulkFormer. PCC: Pearson correlation coefficient. SCC: Spearman correlation coefficient. NES: normalized enrichment score. HR: hazard ratio. Statistical tests: The prognostic markers shown in (h) were identified using univariate Cox regression analysis. Survival curves in (i) were compared using the log-rank test. Significance levels are indicated as follows: *p < 0.05, **p < 0.01, ***p < 0.001.

As BulkFormer imputes masked values based on contextualized expression patterns, its performance is expected to depend on the amount of contextual information available. To test this hypothesis, we evaluated imputation performance across a range of masking ratios on the test set. Consistent with expectations, performance peaked when the test-time masking ratio matched the pretraining condition (15%; r = 0.949) and gradually declined as the masking ratio increased (**Fig. 3b**). Notably, when the masking ratio reached 35%, the performance dropped to r = 0.752, comparable to simple mean- or median-based imputation. These findings suggest that BulkFormer is particularly advantageous when the proportion of missing genes is below 35%, while simpler statistical imputation methods may suffice in cases of more extreme sparsity. To further assess its generalizability, we evaluated BulkFormer on an external test set comprising bulk RNA-seq profiles of 1,000 cancer patients randomly selected from the cancer genome atlas (TCGA) dataset. The model maintained strong imputation performance, achieving a PCC of r = 0.914 (**Fig. 3c**), demonstrating its robustness across diverse biological contexts.

Pancreatic cancer is among the deadliest malignancies, often referred to as the “king of cancers,” and poses a serious threat to human health^30^. Bulk RNA-seq enables the identification of differentially expressed genes (DEGs) between tumor and adjacent normal tissues in patients with pancreatic cancer, offering insights into tumorigenic mechanisms and opportunities for therapeutic development. However, due to technical limitations, the expression of certain potential cancer-driver genes may be missing during sequencing, thereby obscuring critical biological signals. In such cases, BulkFormer’s contextualized imputation capabilities can be leveraged to recover missing gene expression values and enhance transcriptomic completeness **(Fig. 3d)**. To test this application, we retrieved a publicly available pancreatic cancer bulk RNA-seq dataset from the GEO database (GEO accession number: GSE132956) and used the limma R package^31^ to identify DEGs between tumor and normal samples, denoted as DEG_1_. We then applied BulkFormer to impute missing gene expression values across all samples and repeated DEG analysis using the same pipeline to obtain a second set of DEGs, denoted as DEG_2_. By computing the set difference between DEG_2_ and DEG_1_, we identified 408 additional DEGs that emerged only after imputation. Gene set enrichment analysis (GSEA)^32^ of these 408 newly uncovered DEGs revealed significant enrichment in biological processes such as cellular respiration, oxidative phosphorylation, and ATP metabolic process **(Fig. 3e).** Notably, oxidative phosphorylation emerged as one of the most significantly enriched pathways, suggesting a potential role in pancreatic tumor biology (**Fig. 3f**). Although cancer cells in hypoxic tumor microenvironments typically rely on anaerobic glycolysis, oxidative phosphorylation has been reported to be upregulated in pancreatic ductal adenocarcinoma, as well as in leukemias and lymphomas. Targeting this pathway with oxidative phosphorylation inhibitors has thus been proposed as a novel therapeutic strategy^33^. These results demonstrate that BulkFormer not only improves transcriptome completeness through imputation, but also enables the discovery of non-canonical, and potentially overlooked, disease mechanisms and therapeutic targets in cancer.

Next, we evaluated the utility of BulkFormer in discovering novel prognostic biomarkers across multiple cancer types. We selected bulk RNA-seq data from eight cancer cohorts in TCGA, including breast invasive carcinoma (BRCA), kidney renal clear cell carcinoma (KIRC), liver hepatocellular carcinoma (LIHC), lung adenocarcinoma (LUAD), ovarian serous cystadenocarcinoma (OV), pancreatic adenocarcinoma (PAAD), skin cutaneous melanoma (SKCM), and stomach adenocarcinoma (STAD). The number of alive and dead patients in each cohort is shown in **Fig. 3g**. For each cancer type, we first identified genes with extremely high missingness, defined as those absent in more than 95% of patient samples. Such genes are usually1excluded from downstream analyses, as conventional imputation methods (e.g., mean or median imputation) assign uniform values across all patients, eliminating potential prognostic signals. Leveraging BulkFormer’s contextualized imputation capabilities, we recovered the expression values of these highly missing genes. Patients were then stratified into high- and low-expression groups based on the median of the imputed expression values, followed by cox proportional hazards regression to identify statistically significant prognostic markers. In all of the cancer types, we identified previously unrecognized protective and risk-associated genes that robustly stratified patient survival (**Fig. 3h**, **Fig. 3i**). Notably, in KIRC, H4C1 emerged as a strong risk factor: patients with high H4C1 expression level exhibited a 5.2-fold higher mortality rate than those with low expression level. In PAAD, PDE6H was identified as a protective factor: patients with high PDE6H expression level had a 74% lower mortality rate (hazard ratio = 0.26) than those with low expression level (**Fig. 3h**, **Fig. 3i**). These newly discovered biomarkers have not been previously reported in the literature, likely due to the high rate of missing expression data. By recovering such data in a biologically informed manner, BulkFormer facilitates the identification of overlooked yet clinically meaningful prognostic biomarkers, with important implications for precision oncology and downstream clinical investigations.

### BulkFormer enables accurate classification of disease types and cancer subtypes from bulk transcriptomes

Disease classification based on transcriptomic profiles is a critical application in clinical research and precision medicine. To evaluate BulkFormer’s power for disease-type annotation, we obtained disease-associated bulk RNA-seq data for 23 major human diseases from the DiSignAtlas^34^ database (**Supplementary Table 3**). We compared BulkFormer with five existing scLLMs as baselines: Geneformer, GeneCompass, scGPT, scFoundation, and scLong. Specifically, transcriptomic embeddings were extracted using BulkFormer and each baseline model, followed by dimensionality reduction using principal component analysis (PCA) to ensure comparable feature spaces. A random forest classifier was then trained for disease classification, and model performance was evaluated using the weighted F1 score. Ten-fold cross-validation showed that BulkFormer achieved the highest overall classification performance (weighted F1 = 0.939), followed by scGPT (weighted F1 = 0.885) (**Fig. 4a**). We further evaluated model performance across individual disease categories and found that BulkFormer consistently outperformed all baselines across nearly all disease types (**Fig. 4b**), highlighting its superior capability in disease annotation compared to existing scLLMs.

**Fig. 4.**
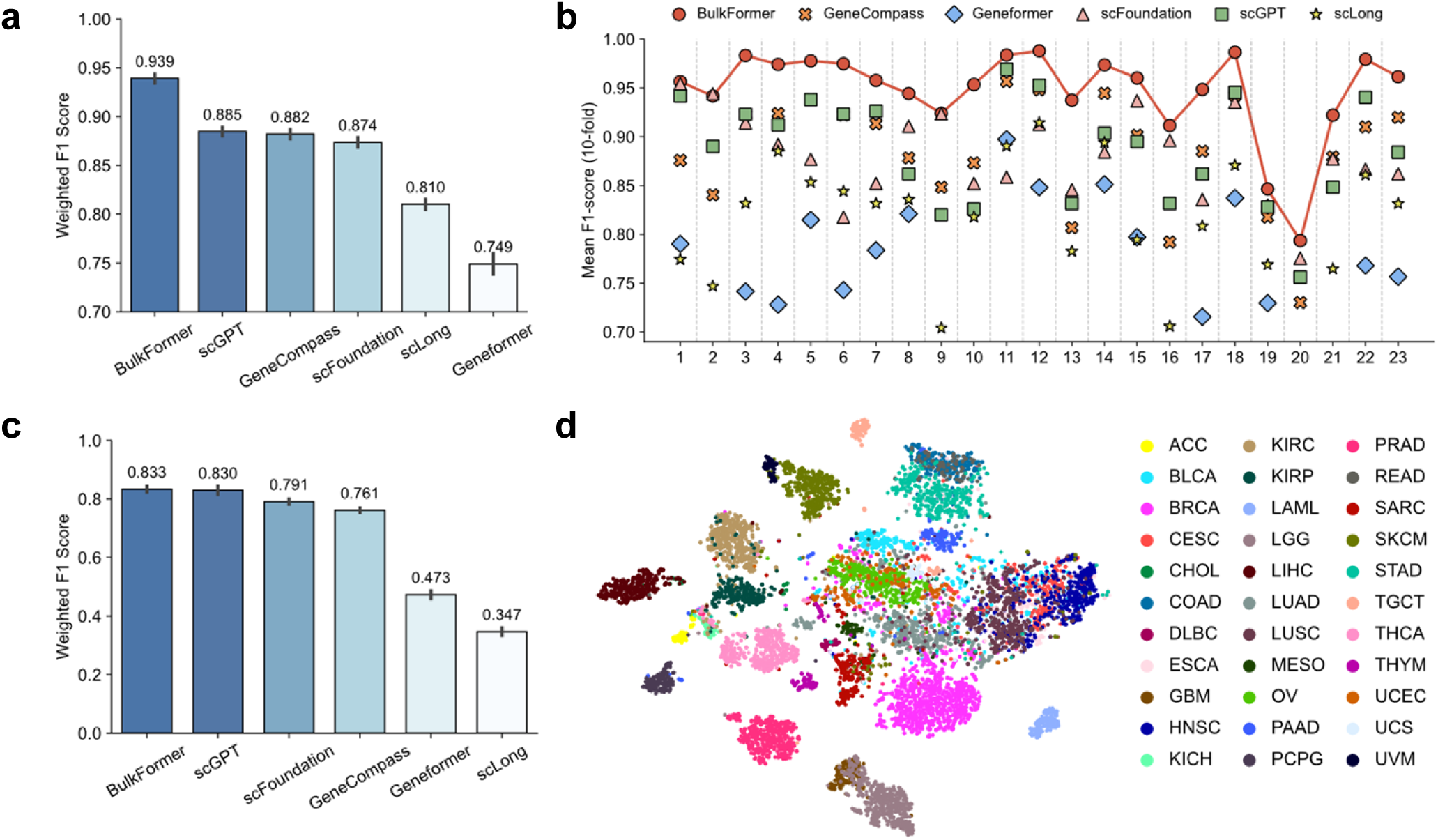
BulkFormer enables accurate classification of disease types and cancer subtypes from bulk transcriptomes. **(a)** Overall performance comparison of BulkFormer and baseline models on the disease classification task. (b) Per-disease classification performance of BulkFormer and other baseline models. (c) Performance comparison on cancer subtype classification across different models. (d) UMAP visualization of sample-level embeddings extracted by BulkFormer for transcriptomes from different cancer types.

We next assessed the ability of BulkFormer to classify cancer subtypes. Bulk RNA-seq data from 33 cancer types were obtained from TCGA (**Supplementary Table 4**), and the same pipeline as in the disease classification task was applied. In this more fine-grained classification task, BulkFormer again achieved the best performance (weighted F1 = 0.833), closely followed by scGPT (weighted F1 = 0.830), while the remaining baseline models performed substantially worse (**Fig. 4c**). To explore the representational quality of BulkFormer embeddings, we visualized the untrained transcriptomic representations of patient samples using uniform manifold approximation and projection (UMAP)^35^. The resulting 2D projections revealed clear clustering by cancer type, with well-separated boundaries between different cancer classes (**Fig. 4d**). We further compared classification performance across each individual cancer type, and observed that BulkFormer and scGPT consistently delivered top-tier performance across nearly all cancer types (**Supplementary Fig. 3**). Together, these results demonstrate that BulkFormer not only enables accurate classification of broad disease categories, but also performs well in more granular subtype classification tasks, such as distinguishing between diverse cancer types.

### BulkFormer enhances prognosis prediction by generating context-aware gene representations

It is an important task to predict the clinical outcomes of cancer patients (survival or death), which are known to be closely linked to their transcriptomic profiles, with the expression levels of certain key genes serving as valuable prognostic biomarkers^36^. To evaluate whether BulkFormer can effectively capture such prognostic signals, we first compared its performance with that of baseline models in predicting patient outcomes based on bulk RNA-seq data.

We collected transcriptomic profiles and survival outcome information for ∼10,000 patients across 33 cancer types from the TCGA database. The ratio of alive to dead patients varied substantially across different cancers (**Fig. 5a**). For each model, we extracted sample-level transcriptomic embeddings, applied PCA to project them into a unified feature space, and then trained a random forest classifier to predict patient survival status. Ten-fold cross-validation revealed that BulkFormer achieved the best performance, with an AUROC of 0.747 and an AUPRC of 0.549. The next-best performer, scFoundation, yielded an AUROC of 0.726 and an AUPRC of 0.520 (**Fig. 5b, 5c**).

**Fig. 5.**
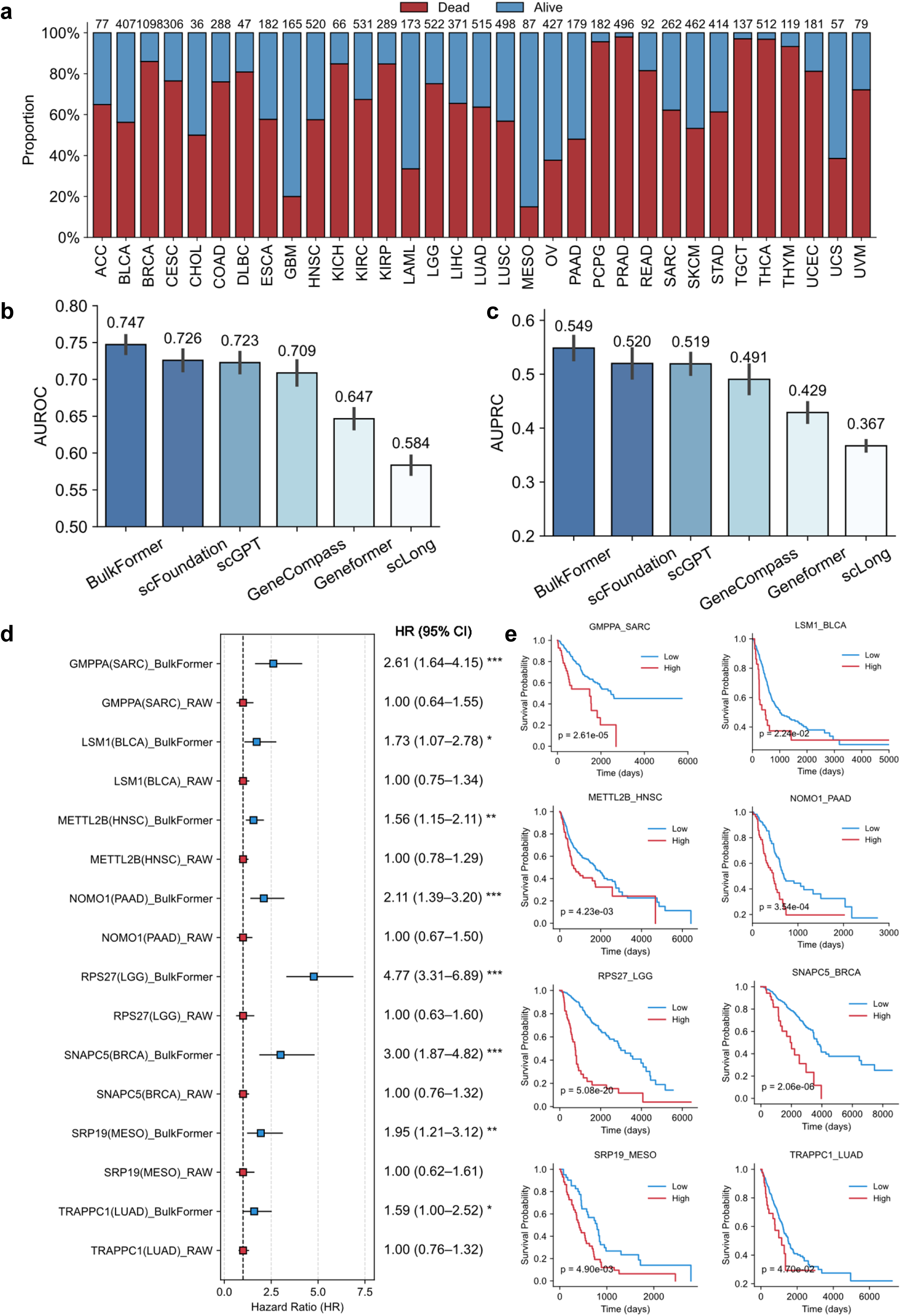
BulkFormer enhances prognosis prediction by generating context-aware gene representations. **(a)** Distribution of survival outcomes (alive vs. dead) across 33 cancer cohorts in the TCGA database. **(b-c)** Performance comparison of BulkFormer and baseline models in predicting patient prognosis. **(d)** BulkFormer-enhanced biomarkers for prognosis prediction. **(e)** Kaplan-Meier survival curves based on BulkFormer-enhanced prognostic biomarkers. AUROC: area under the receiver operating characteristic curve. AUPRC: area under the precision-recall curve. HR: hazard ratio. Statistical tests: The prognostic markers shown in (d) were identified using univariate Cox regression analysis. Survival curves in (e) were compared using the log-rank test. Significance levels are indicated as follows: *p < 0.05, **p < 0.01, ***p < 0.001.

While BulkFormer outperformed all baselines, there remains considerable room for improvement. A key challenge in prognosis modeling lies in the high level of noise in bulk transcriptomic data, because many genes may be unrelated to patient outcomes. To mitigate this, conventional approaches often rely on univariate cox regression to identify prognostically relevant genes, and subsequently combine their expression values to compute risk scores. However, this strategy may overlook weakly expressed or subtly variable genes that nonetheless carry latent prognostic information. To address this limitation, we leveraged BulkFormer’s context-aware gene embeddings to amplify the predictive power of otherwise insignificant genes. Specifically, we adopted a supervised learning framework in which BulkFormer was used to extract contextualized gene embeddings for each patient sample. A random forest classifier was then trained to predict survival probability using these embeddings. Using 10-fold cross-validation, we obtained model-derived prediction scores for each patient, which were subsequently used to stratify patients into high- and low-risk groups based on the median score. Cox and log-rank tests were then applied to assess the prognostic value of these new model-derived scores. We applied this strategy across eight representative cancer types and demonstrated enhanced prognostic resolution for genes that were initially non-significant. Remarkably, RPS27, a gene that originally showed no prognostic value in lower-grade glioma (LGG; HR = 1.0), became a highly significant risk factor after contextual embedding by BulkFormer (HR = 4.77) (**Fig. 5d, 5e**). These results suggest that BulkFormer-generated gene embeddings can rescue weak or overlooked biomarkers and enhance their prognostic utility, providing new opportunities for clinical research and precision medicine.

### BulkFormer facilitates *in silico* drug discovery by modeling compound–transcriptome relationships

Transcriptomic profiles are essential for *in silico* drug discovery, serving as both representations of cellular identity and as response signatures following drug perturbation. Compounds with similar mechanisms of action typically induce concordant transcriptomic changes, whereas pharmacologically distinct agents elicit divergent expression profiles. Identifying compounds that induce transcriptomic responses inversely correlated with disease-associated signatures enables phenotype-based drug screening. Furthermore, functional enrichment of drug-induced gene expression changes can provide mechanistic insight into compound activity^37^. PRnet is a deep learning framework designed to predict transcriptomic responses to compound perturbations, trained on data from the LINCS^38^ database. The LINCS database provides a large-scale resource of gene expression profiles from various compounds and cell lines under perturbation conditions. PRnet employs a denoising autoencoder to integrate untreated transcriptomic features with compound representations, which are then decoded into predicted post-treatment expression profiles^39^. However, PRnet reduces raw gene expression profiles via linear layers before integration, treating the transcriptome at a coarse resolution and overlooking gene-specific drug effects. To address this limitation, we incorporated context-aware gene embeddings produced by BulkFormer and scLLMs, fusing them with compound features at the gene level for more fine-grained representation. These fused features were passed through an MLP to predict the post-treatment expression levels of individual genes. We used the same dataset and data-splitting protocol as PRnet and evaluated model performance using PCC and spearman correlation coefficients (SCC) between predicted and ground-truth transcriptomic responses. Our results demonstrate that models using gene-level fusion consistently outperformed PRnet, indicating the effectiveness of this strategy. Notably, the model trained with BulkFormer-generated gene embeddings achieved the best performance (PCC = 0.493; SCC = 0.430), substantially outperforming PRnet (PCC = 0.408; SCC = 0.345) and scLLM-based baselines (**Fig. 6a, 6b**). To assess the biological relevance of the predicted gene expression responses, we evaluated the case of Dovitinib **(Fig. 6c**), an orally bioavailable, potent inhibitor of class III–V receptor tyrosine kinases, which was not present in the training set. BulkFormer accurately predicted the Dovitinib-induced transcriptomic changes, which were subsequently analyzed via GSEA using the KEGG pathway database. The results revealed significant downregulation of cancer-related pathways, including colorectal cancer, pancreatic cancer, and breast cancer (**Fig. 6d, 6e**). Consistent with these enrichment results, previous studies have shown that Dovitinib combined with oxaliplatin exhibits enhanced in vitro cytotoxicity in colon cancer cell lines, regardless of RAS-RAF status^40^.

**Fig. 6.**
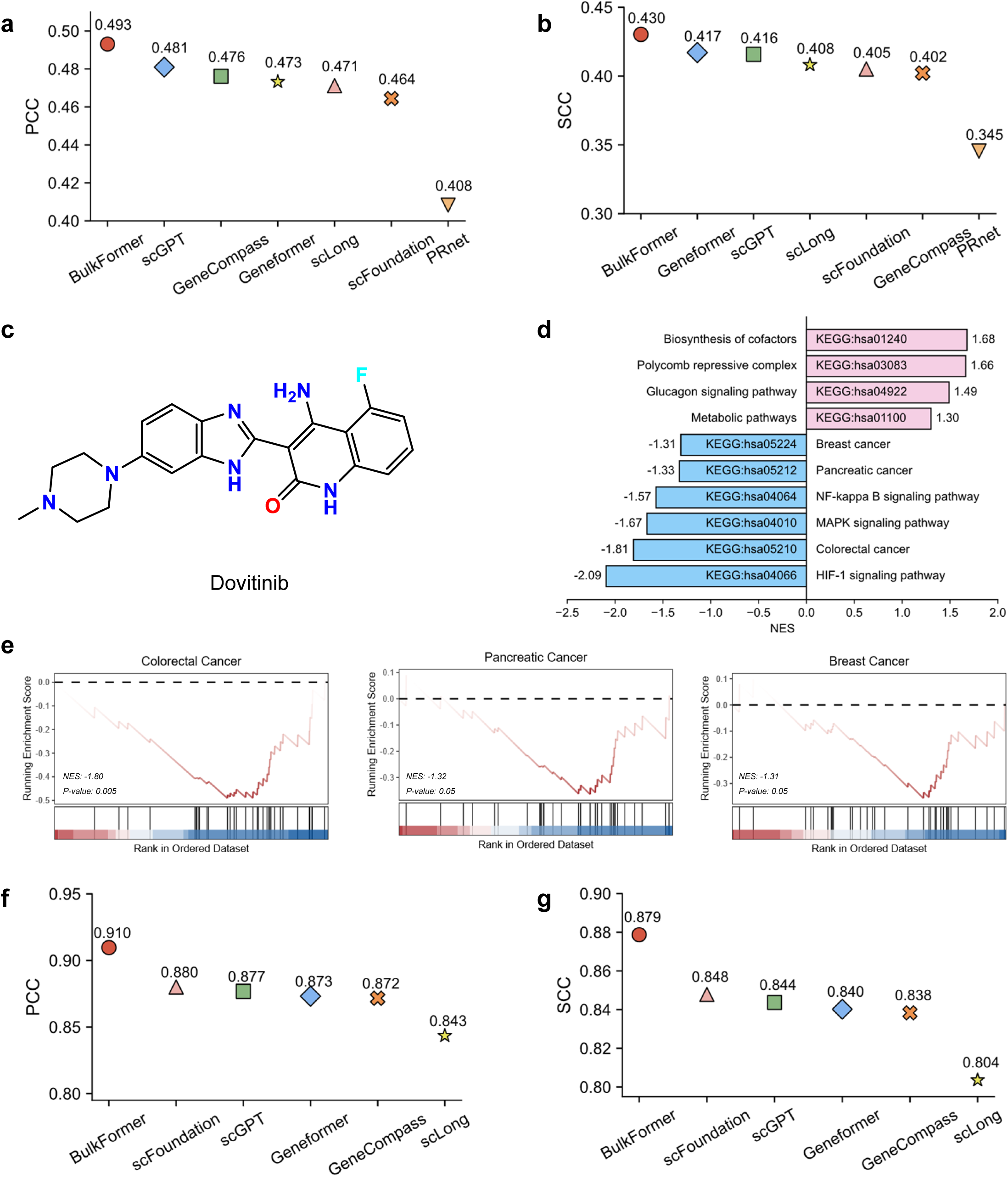
BulkFormer facilitates *in silico* drug discovery by modeling compound–transcriptome relationships. **(a-b)** Performance comparison of BulkFormer and baseline models on the compound perturbation prediction task. **(c)** Chemical structure of the compound Dovitinib. **(d-e)** GSEA enrichment analysis results based on differentially expressed genes predicted by BulkFormer following Dovitinib perturbation. **(f-g)** Performance comparison of BulkFormer and baseline models on the drug response prediction task. PCC: Pearson correlation coefficient. SCC: Spearman correlation coefficient. NES: normalized enrichment score.

Drug response prediction is another crucial component of *in silico* drug discovery, aiding in the identification of compounds with selective sensitivity or resistance across cancer types^41^. To evaluate this task, we evaluated BulkFormer and scLLMs on a large-scale drug response dataset curated from the Genomics of Drug Sensitivity in Cancer (GDSC)^42^ database, which includes IC50 values for 255 compounds across 700 cancer cell lines representing 32 cancer types. Compound features were extracted using the pretrained compound language model KPGT^43^, while transcriptomic representations of cell lines were derived from BulkFormer and baseline models. The features were concatenated and passed through an MLP to predict IC50 values. Ten-fold cross-validation demonstrated that BulkFormer again achieved the best performance (PCC = 0.910; SCC = 0.879), followed by scFoundation (PCC = 0.880; SCC = 0.848) (**Fig. 6f, 6g**), confirming its utility for predictive modeling of drug sensitivity.

### BulkFormer improves gene essentiality prediction based on bulk transcriptomic profiles

Gene essentiality refers to the extent to which a gene is critical for the survival and development of an organism. Compared to non-essential genes, disruption of essential genes often leads to severe consequences, including lethality^44^. In cancer cells, gene essentiality scores are typically defined as the impact of gene knockout on cellular viability. Thus, targeting essential genes that sustain cancer cell growth represents a promising therapeutic strategy^45^. Gene essentiality is influenced by various biological factors, with gene expression levels being among the most important. However, existing methods rarely enable direct prediction of gene essentiality from expression profiles alone. To address this gap, we compiled a dataset from the DepMap^46^ database, consisting of gene expression levels and gene essentiality scores for 17,862 protein-coding genes across 1,103 human cancer cell lines. We then applied BulkFormer and scLLMs to generate context-aware gene embeddings from the gene expression profiles. These embeddings were used as input to an MLP to predict the essentiality score for each gene. Ten-fold cross-validation results showed that BulkFormer significantly outperformed all baseline models, achieving a PCC of 0.931 and a SCC of 0.759 (**Supplementary Fig. 4**). These results demonstrate that BulkFormer can more accurately infer the essentiality of individual genes based solely on transcriptomic data, enabling rapid identification of cancer-specific vulnerabilities from patient-derived expression profiles. This capability has important implications for precision oncology, offering a potential strategy to prioritize therapeutic targets for cancer treatment.

## Discussion

Recent advances in scLLMs have significantly improved the modeling of sparse scRNA-seq data and achieved state-of-the-art performance across a range of single-cell tasks. It is known that scRNA-seq typically detects only ∼3000 genes per cell, while bulk RNA-seq can profiling ∼16,000 genes per sample. Due to this fundamental difference in data modality, however, scLLMs pretrained on sparse single-cell data often underperform on clinical and tissue-level tasks that rely on bulk RNA-seq. This limitation motivated the development of a dedicated foundation model specifically designed for bulk transcriptomic data.

In this study, we developed BulkFormer, a large-scale foundation model specifically designed for bulk-level transcriptomic modeling. BulkFormer employs a hybrid encoder architecture that integrates a GNN to capture explicit gene–gene relationships with a performer module to learn implicit expression dependencies. This structure enables the model to capture both biological priors and context-dependent gene interactions. BulkFormer was pretrained on over 500,000 bulk transcriptomic profiles, encompassing the expression of approximately 20,000 protein-coding genes, allowing it to learn comprehensive and biologically meaningful representations.

Across a series of bulk transcriptome-related downstream tasks, including transcriptome imputation, disease annotation, prognosis modeling, drug response prediction, compound perturbation prediction, and gene essentiality prediction, BulkFormer consistently outperformed existing baseline and scRNA-seq foundation models while incurring substantially lower training costs. Notably, its high-quality context-aware gene embeddings and superior imputation performance enabled the discovery of previously unrecognized prognostic biomarkers and the enhancement of weakly predictive known markers, providing valuable insights for cancer prognosis and biomarker development.

Despite its strong performance, BulkFormer still face some limitations. Its performance on single-cell specific tasks such as cell type annotation is limited compared to scLLMs because it was pretrained on bulk RNA-seq data. This discrepancy reflects the intrinsic modality gap between bulk and single-cell transcriptomics. As such, we recommend that users select foundation models based on their purpose to ensure optimal performance. In addition, BulkFormer focuses exclusively on protein-coding genes, and does not include non-coding transcripts, which restricts its utility for applications involving non-coding RNA biology.

In the future, BulkFormer opens several avenues for future research. First, the development of a unified RNA-seq foundation model capable of jointly modeling bulk and single-cell data could bridge modality gaps and leverage the complementary strengths of both platforms. Second, expanding the model to include non-coding genes may provide new opportunities for studying regulatory RNA biology. Finally, current transcriptome foundation models, including BulkFormer, primarily rely on gene expression matrices derived from RNA-seq data, while largely overlooking sample-level meta-information. Such metadata, including variables like sex, age, disease type, and tissue of origin, may contain critical covariates that influence gene expression patterns. Effectively integrating these metadata with transcriptomic profiles represents an important future direction in the development of next-generation transcriptome foundation models.

## Methods

### Construction of the PreBULK dataset

#### Data collection

Unlike scRNA-seq data, there is a notable lack of centralized public resources that systematically curate bulk RNA-seq datasets. To address this, we manually collected human bulk RNA-seq profiles from public repositories including the Gene Expression Omnibus (GEO) and ARCHS4. Duplicate entries were removed based on GEO sample identifiers. To ensure data consistency and compatibility for model pretraining, only samples with available raw count matrices were retained. The resulting dataset spans nine major human physiological systems and includes samples from both healthy and diseased states.

#### Gene ID unification

To ensure consistency across datasets, all gene expression matrices were unified using Ensembl gene identifiers, as provided by the Ensembl^47^ database. We included all annotated human protein-coding genes, totaling 20,010 genes. For samples in which specific gene entries were absent, zero-padding was applied to maintain a consistent dimensionality across all expression profiles.

#### Quality control

To remove low-quality samples, we filtered the dataset using the NumPy package by excluding expression profiles with fewer than 14,000 non-zero gene values. This threshold effectively eliminates potentially misclassified or contaminated scRNA-seq data, reduces sparsity, and ensures the retained samples reflect true bulk transcriptomic profiles. Finally, the constructed PreBULK dataset comprises a total of 522,769 bulk transcriptomic samples.

#### Data preprocessing

To correct for differences in sequencing depth and gene length, and to scale raw count values to a comparable range, we converted raw gene expression counts to transcripts per million (TPM). The TPM value for each gene was calculated as follows:

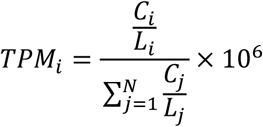

Where *C*_*i*_ is the raw count of gene *i*, *L*_*i*_ is the length of gene *i* and *N* is the total number of genes. This normalization ensures that gene expression levels are comparable both within and across samples.

### BulkFormer model architecture

#### Embedding module

The core principle of LLM-like models is to represent tokens as high-dimensional embeddings, which are optimized through pretraining such that semantically related tokens acquire similar representations. In the context of transcriptomic data, this involves two key components: the representation of gene identities and the encoding of gene expression values.

To represent gene identities, we retrieved the primary protein product sequence of each gene from the Ensembl database and extracted sequence-based embeddings using the ESM2 model. These protein embeddings were aggregated using mean pooling to generate the initial gene embedding. Compared with one-hot encoding, ESM2–based embeddings capture functional and evolutionary relationships among genes more effectively, thereby enhancing BulkFormer’s ability to learn biologically meaningful gene–gene dependencies.

Gene expression values within a transcriptome can be interpreted as defining an internal ordering of gene tokens, reflecting their relative biological activity. A naïve approach would involve rank-based encoding, but this method suffers from two major limitations. First, it reduces data resolution by discarding the absolute magnitude of expression values. Second, rank values are sample-specific and not directly comparable across samples. For example, two genes with the same rank in different samples may have substantially different expression magnitudes. To overcome these limitations, we propose a rotary expression embedding (REE) strategy inspired by rotary position encoding (ROPE). While ROPE is traditionally used to encode positional information in natural language processing, we adapt this concept to embed absolute gene expression values, preserving both their magnitude and continuity. Specifically, for gene *g* with expression value *x*_*g*_, we defined the expression embedding as:

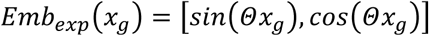

where the frequency matrix is defined as:

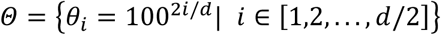

and *d* is the embedding dimension. Prior to this transformation, expression values are normalized using *log*(*TPM* + 1) to ensure numerical stability.

In addition, we incorporated sample-level context by applying an MLP to compress the entire gene expression vector of each sample into a low-dimensional embedding, capturing its global transcriptomic state.

Finally, the three types of embedding, including the gene identity embedding *E*_*ESM*2_, the expression value embedding *E*_*REE*_, and the sample context embedding *E*_*MLP*_, are combined through element-wise addition to generate the final model input.

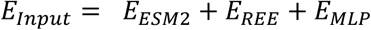

### BulkFormer blocks

The core architecture of BulkFormer consists of *N* stacked BulkFormer blocks, each comprising a graph convolutional network (GCN) layer followed by *K* layers of performer-based self-attention. The GCN layers are designed to capture explicit gene–gene relationships derived from biological prior knowledge, while the Performer layers model global, implicit dependencies across genes.

To incorporate prior biological knowledge, we first construct a gene-gene relationship graph *G* = (*V*, *E*), where *V* is the set of genes and *E* denotes the edges representing gene-gene associations derived from known gene knowledge graph. In the ablation studies, we systematically evaluated the impact of different knowledge graphs on model performance, including the gene co-expression graph, the gene ontology (GO) similarity graph, and the protein–protein interaction (PPI) graph. Among these, the gene co-expression graph achieved the best performance and was therefore adopted in the GCN module of BulkFormer (**Supplementary Fig. 5**). We constructed the gene co-expression graph using the absolute values of Pearson correlation coefficients (PCC) between gene expression profiles as edge weights. To avoid excessive graph density and retain the most informative interactions, we preserved only the top 20 edges with the highest weights for each node. Additionally, edges with PCC below 0.4 were discarded to eliminate weak or spurious gene–gene associations.

The GCN layers updates the gene embeddings based on the graph structure as follows:

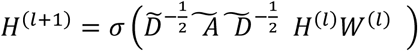

where, *H*^(*l*)^ denotes the feature matrix at the *l* − *th* layer, 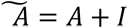 is the adjacency matrix with self-loops, 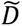 is the corresponding degree matrix, *W*^(*l*)^ is a learnable weight matrix, and *σ*(·) is a nonlinear activation function.

Following the GCN layers, we employed Performer, a kernel-based transformer variant, to capture long-range gene-gene interactions in transcriptomic profiles. Unlike standard self-attention mechanisms, Performer approximates attention scores via a kernelizable formulation to achieve linear scalability. Specifically, the attention computation is expressed as:

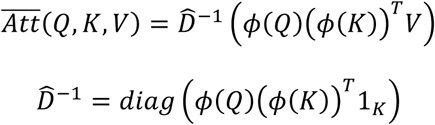

where *Q* = *XW*_*q*_, *K* = *XW*_*k*_ and *V* = *XW*_*v*_ are linear transformation of the input *X*, *W* are training parameters, *ϕ*(·) is a kernel function that used for approximating the exponential attention matrix, 1_*K*_ is the all-ones vector of length *K*, and *diag*(·) is a diagonal matrix with the input vector as the diagonal. This formulation eliminates the quadratic dependency on sequence length, enabling efficient modeling of large-scale transcriptomic inputs while preserving key interaction patterns.

After passing through the stacked BulkFormer blocks, the updated gene representations are then fed into an MLP to output the final gene expression values.

### BulkFormer pretraining task

During the pretraining stage, BulkFormer was trained using a masked language modeling (MLM) strategy in a self-supervised manner. Specifically, approximately 15% of gene expression values in each input transcriptome were randomly selected and masked using a special placeholder token (−10), resulting in a partially masked input vector. The model is then tasked with reconstructing the masked expression values based on the unmasked context. The training objective is to minimize the reconstruction error between the predicted and true expression values at the masked positions, using the mean squared error (MSE) as the loss function ℒ:

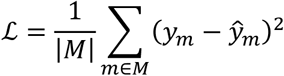

where *M* denotes the set of masked gene indices, *y*_*m*_ is the true expression value at position *m*, and *ŷ_m_* is the model’s corresponding predicted value.

All parameters 𝛩 = {*E*_*Input*_, 𝛩_*GCN*_, 𝛩_*Performer*_, 𝛩_*MLP*_} were optimized during the pretraining stage. The detailed hyperparameter setting of the BulkFormer can be found in **Supplementary Table 5**.

### BulkFormer implementation details

BulkFormer was implemented in PyTorch (v2.5.1) with CUDA 12.4 and Python 3.12.7. Pretraining was conducted on eight NVIDIA A800 GPUs for 29 epochs, requiring approximately 350 GPU hours. We used the AdamW optimizer with a max learning rate of 0.0001. The learning rate was linearly warmed up from zero, reaching its peak after 5% of total training steps. To support large model training, we adopted a per-device batch size of 4 and employed gradient accumulation with 128 steps, resulting in a large effective batch size.

### Downstream methods

BulkFormer’s downstream tasks fall into three major categories: (1) Expression-level prediction tasks: These involve imputing missing gene expression values in bulk transcriptomes using context-aware predictions directly output by BulkFormer. (2) Gene-level embedding tasks: These leverage gene embeddings produced by the final Performer layer of BulkFormer as input features for downstream applications such as compound perturbation prediction, gene essentiality estimation, and prognostic modeling of individual genes. (3) Sample-level embedding tasks: In these tasks, gene embeddings from the final Performer layer are aggregated via max pooling to obtain sample-level representations. These representations are then used for downstream analyses, including disease classification, cancer subtype annotation, drug response prediction, and patient prognosis modeling.

We compared BulkFormer against five representative scLLMs, including Geneformer, GeneCompass, scGPT, scFoundation, and scLong. Due to differences in model architectures and pretraining objectives, we adopted task-specific evaluation strategies for fair comparison. For expression-level prediction tasks, only scFoundation and scLong were included as baselines, since they are the only models pretrained to reconstruct gene expression values directly. For gene-level embedding tasks, we excluded scLong and focused on the remaining scLLMs. As these models were pretrained using only the top 1,000–2,000 highly expressed genes per cell rather than full-length transcriptomes, we divided each bulk transcriptome into ten non-overlapping blocks and sequentially input them into the scLLMs to extract gene-level embeddings. For sample-level embedding tasks, we used the top 2,000 most highly expressed genes from each bulk sample and applied max pooling over their embeddings to generate sample-level representations for downstream classification and regression tasks.

### Evaluation metrics

We used several quantitative metrics to evaluate the performance of BulkFormer and baseline models across different downstream tasks.

### Pearson correlation coefficient (PCC)

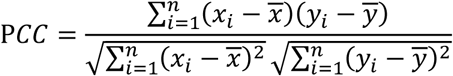

### Spearman correlation coefficient (SCC)

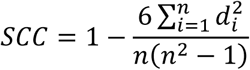

where *d*_*i*_ is the difference between the ranks of corresponding variables.

### Area under the receiver operating characteristic curve (AUROC)

AUROC evaluates the ability of a model to distinguish between positive and negative classes across different decision thresholds. It is computed as the area under the ROC curve.

### Area under the precision-recall curve (AUPRC)

AUPRC summarizes the trade-off between precision and recall. Particularly useful for imbalanced datasets, it is calculated as the area under the precision-recall curve.

### F1 score

The F1 score is the harmonic mean of precision and recall, defined as:

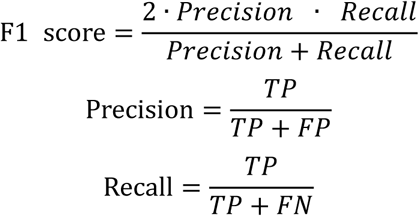

where TP (True Positive) denotes the number of samples correctly predicted as the positive class, FP (False Positive) represents the number of negative samples incorrectly predicted as positive, and FN (False Negative) refers to the number of positive samples incorrectly predicted as negative.

### Weighted F1 score

For multi-class settings, the weighted F1 is defined as:

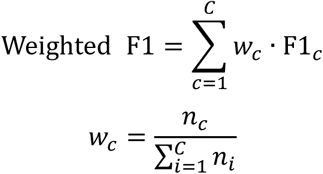

Where *C* is the number of classes, *n*_*c*_ is the number of samples in class *c*, and F1_*c*_ is the F1 score for class *c*.

## Data availability

The large-scale human bulk transcriptomes were collected from the GEO (https://www.ncbi.nlm.nih.gov/geo/) and the ARCHS4 (https://archs4.org/) database. The transcriptomic profiles and corresponding survival information of cancer patients were downloaded from the TCGA (https://portal.gdc.cancer.gov/) database. The data used for the disease classification task were obtained from the DiSignAtlas (http://www.inbirg.com/disignatlas/home) database. Compound perturbation data were obtained from the LINCS (https://clue.io/) database. Drug response data were accessed from the GDSC (https://www.cancerrxgene.org/) database. Gene essentiality data were downloaded from the DepMap (https://depmap.org/portal/) database. The data used in this study are available on Zenodo (https://doi.org/10.5281/zenodo.15559368). Source data are provided with this paper.

## Code availability

BulkFormer source code is available on GitHub (https://github.com/KangBoming/BulkFormer).

## Acknowledgements

This study was supported by the grants from the National Natural Science Foundation of China [62025102].

## Author information

These authors contributed equally: Boming Kang, Rui Fan.

## Authors and Affiliations

**Department of Biomedical Informatics, State Key Laboratory of Vascular Homeostasis and Remodeling, School of Basic Medical Sciences, Peking University, 38 Xueyuan Rd, Beijing, 100191, China**

Boming Kang, Rui Fan & Qinghua Cui

**School of Pharmaceutical Sciences, Jilin University, Changchun 130021, China**

Meizheng Yi

**School of Sports Medicine, Wuhan Sports University, No. 461 Luoyu Rd. Wuchang District, Wuhan 430079, Hubei Province, China**

Chunmei Cui & Qinghua Cui

## Contributions

BK and RF designed the study. BK, RF and MY performed the study. BK, RF, MY, CC and QC wrote or edited the manuscript. QC supervised the study.

## Corresponding author

Correspondence to Qinghua Cui

## Supplementary Figures

**Supplementary Figure 1.**
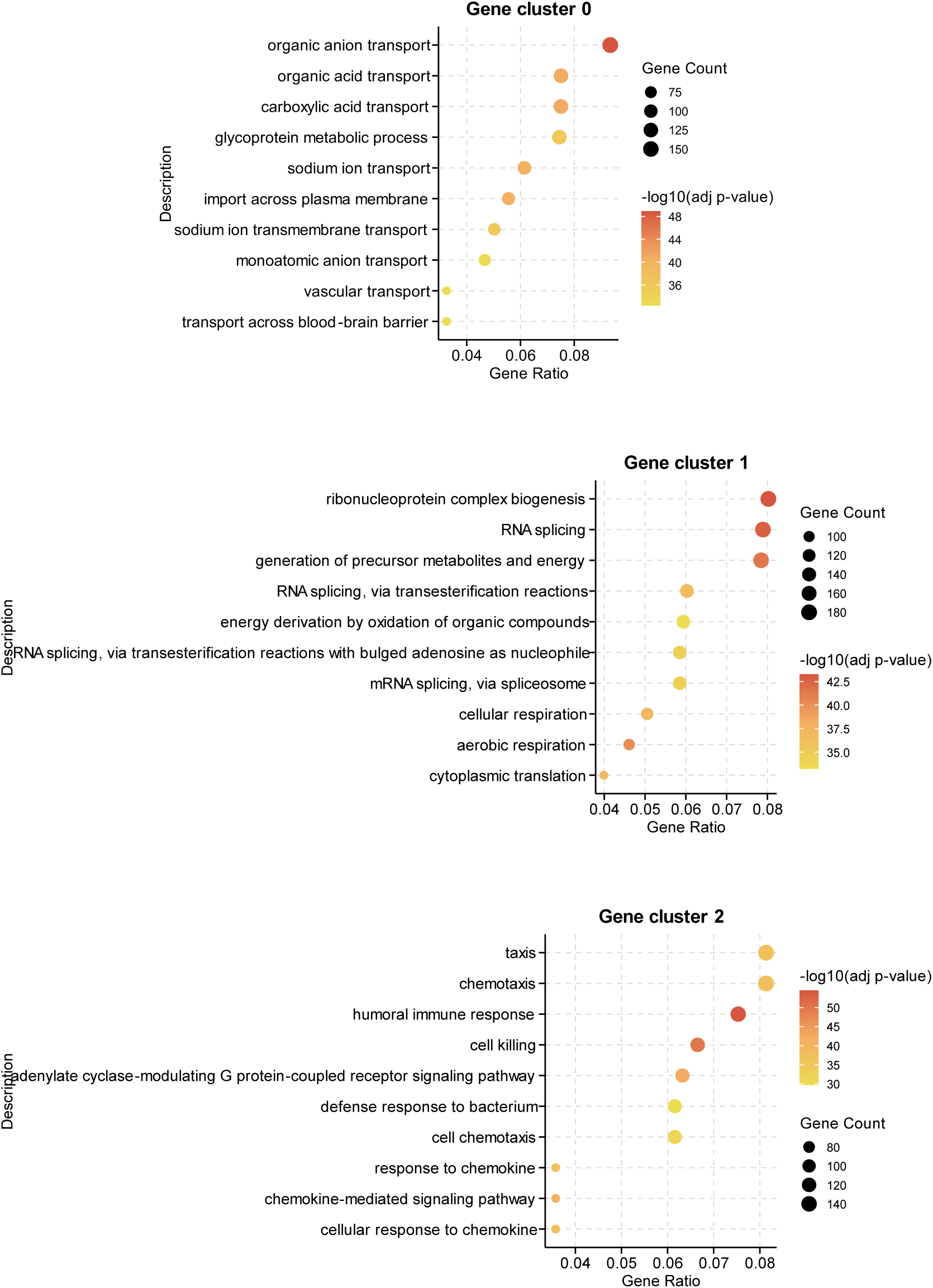
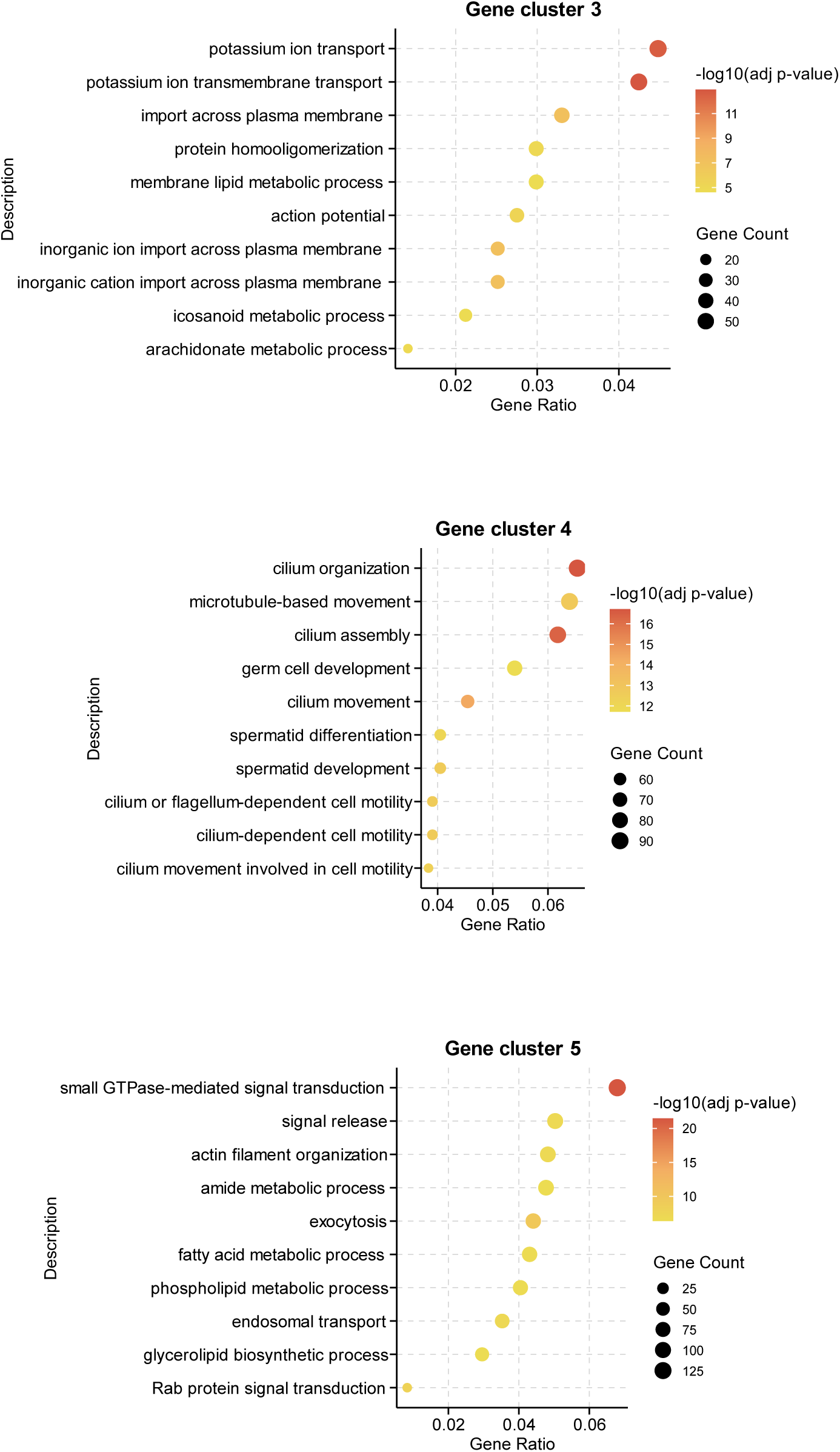
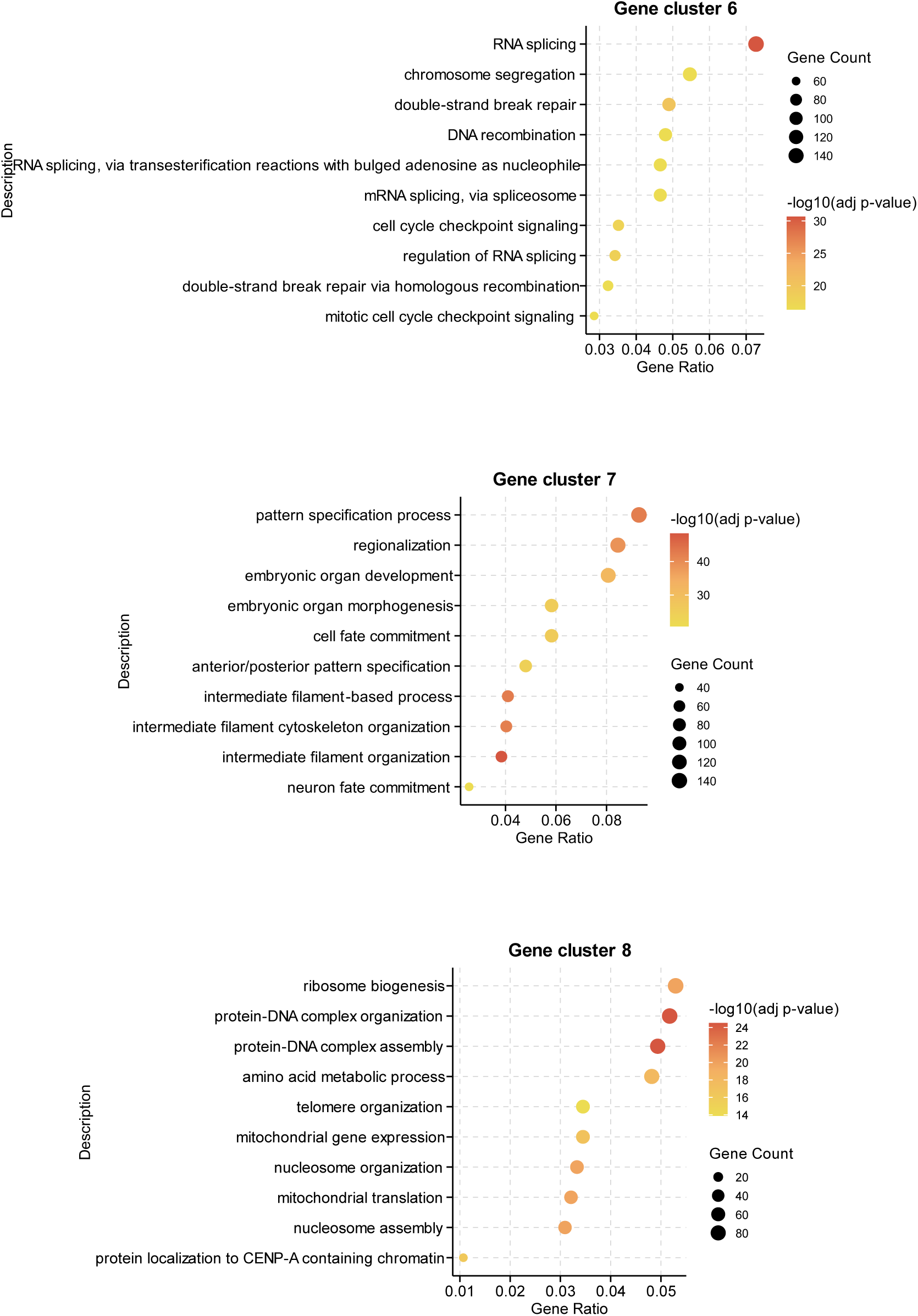
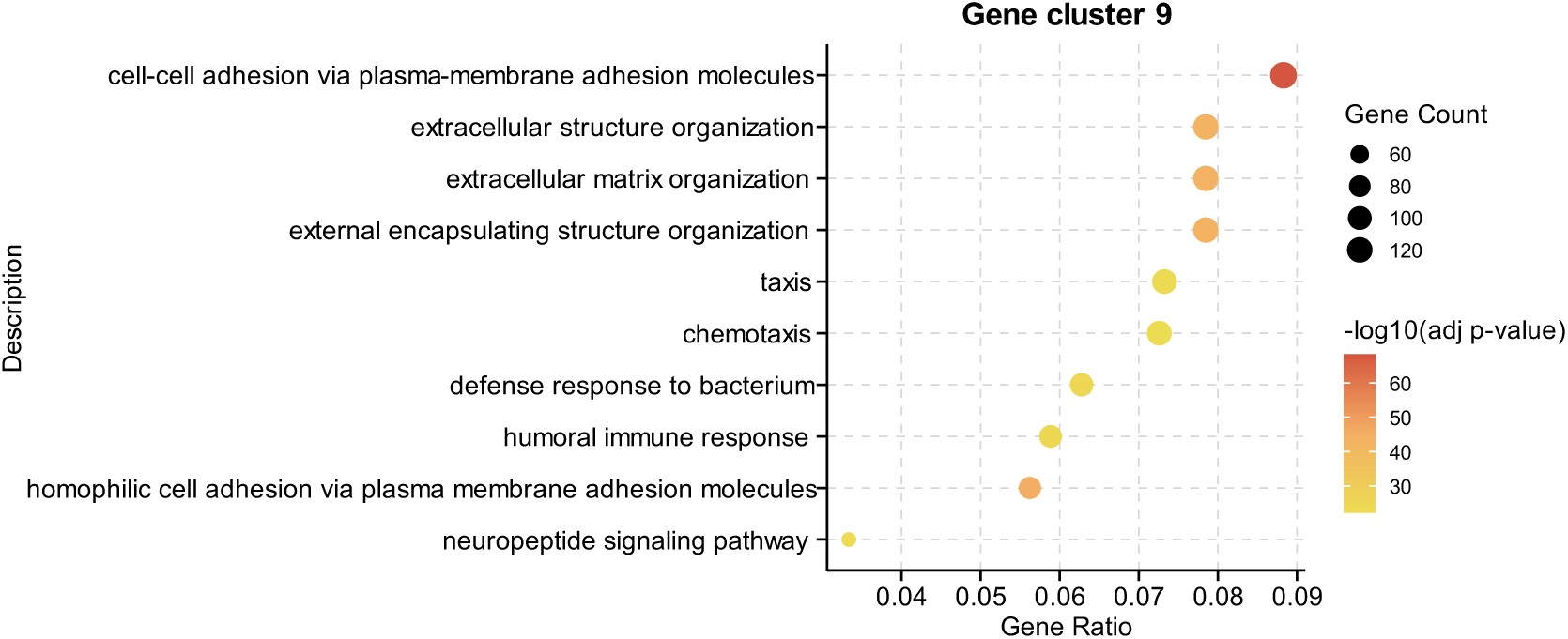
GO enrichment analysis of distinct gene clusters derived from BulkFormer-generated gene embeddings.

**Supplementary Figure 2.**
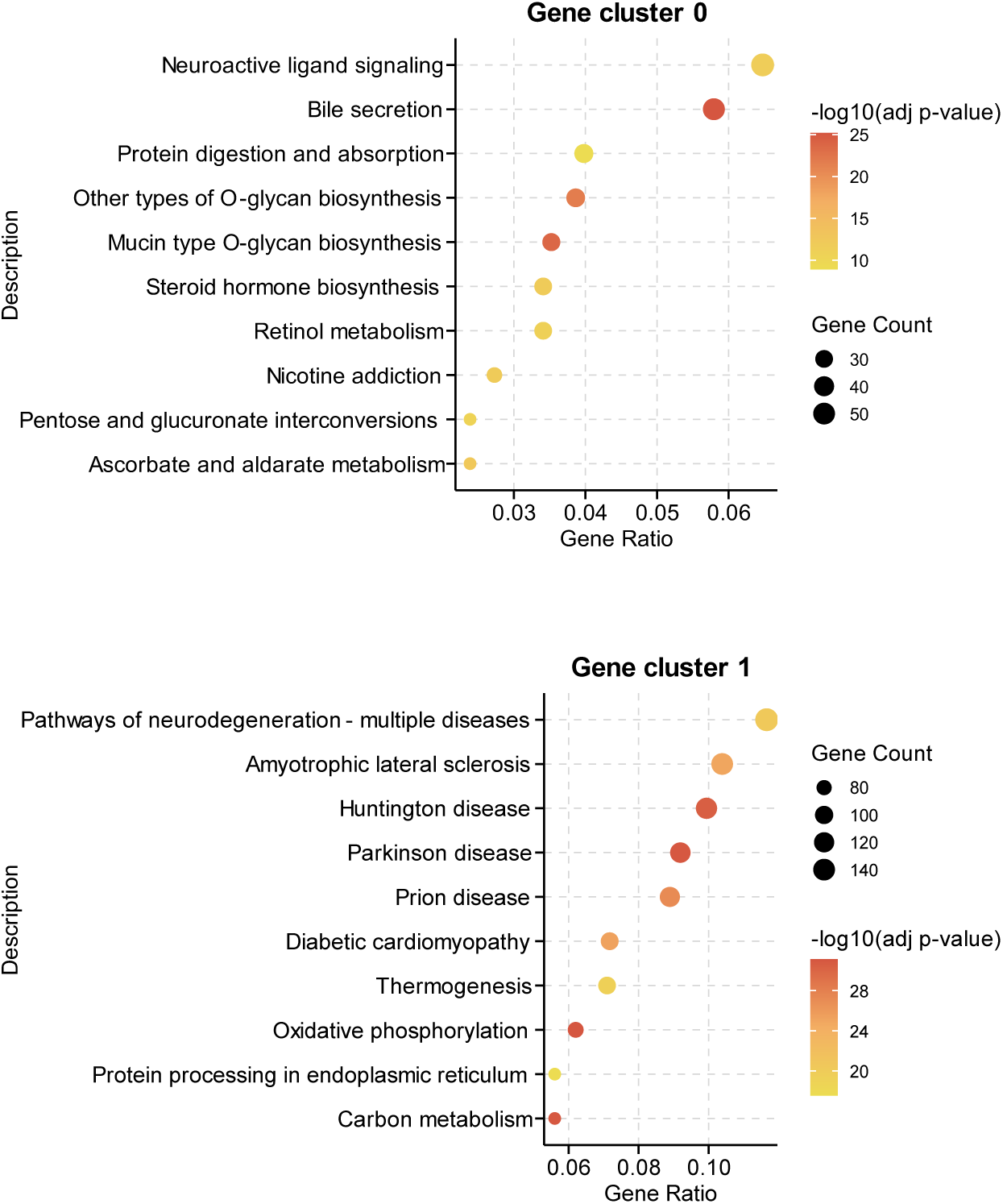
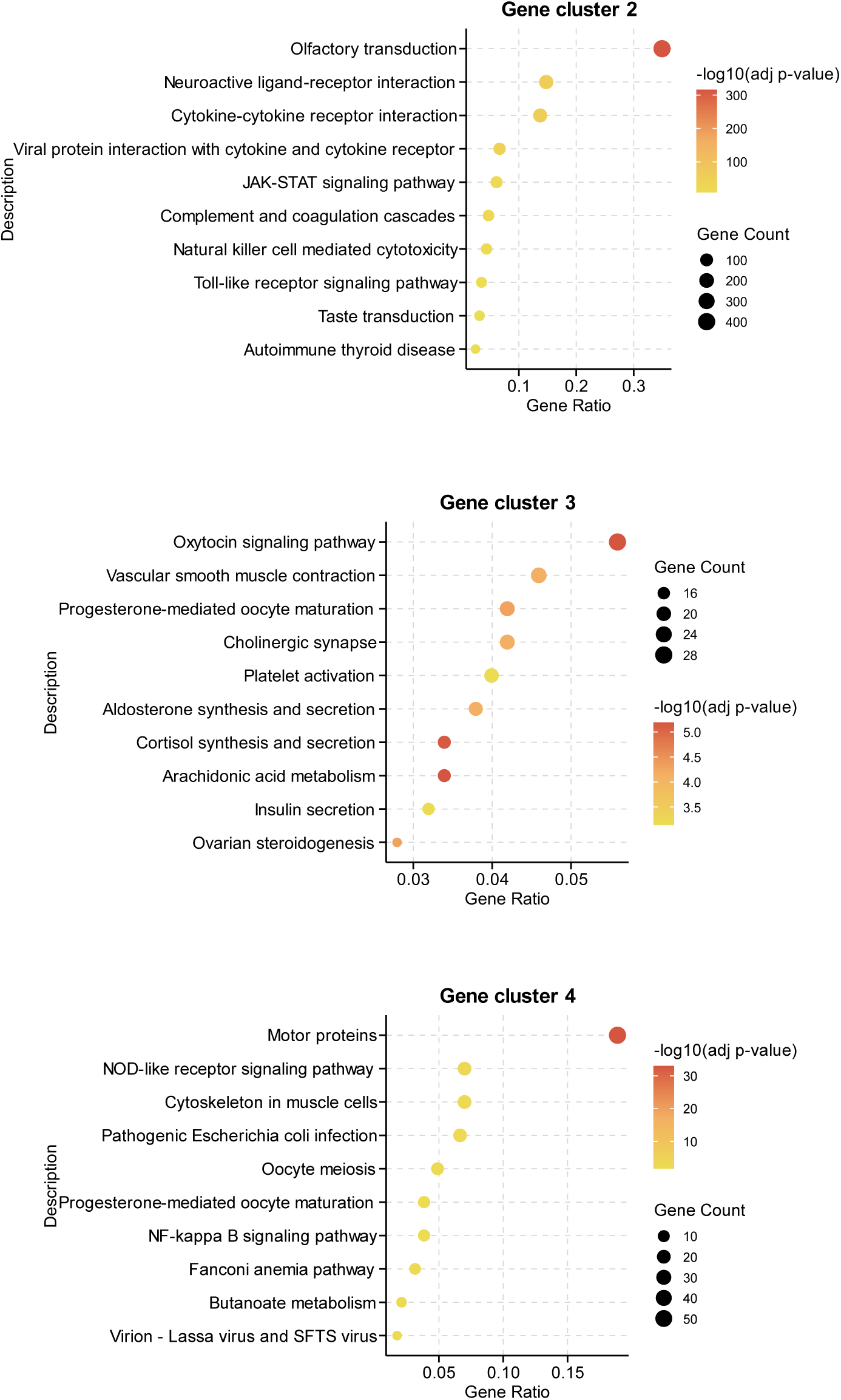
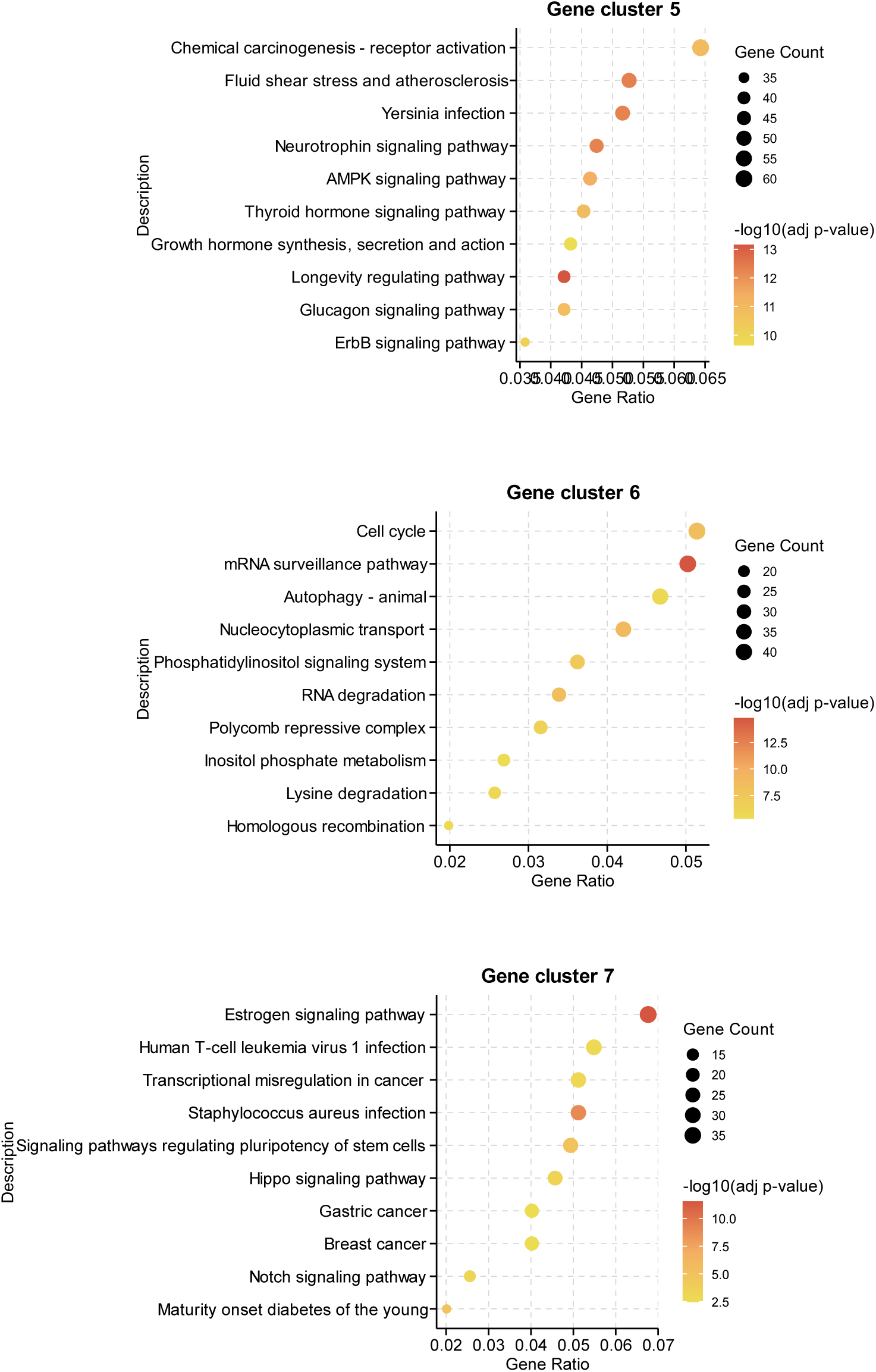
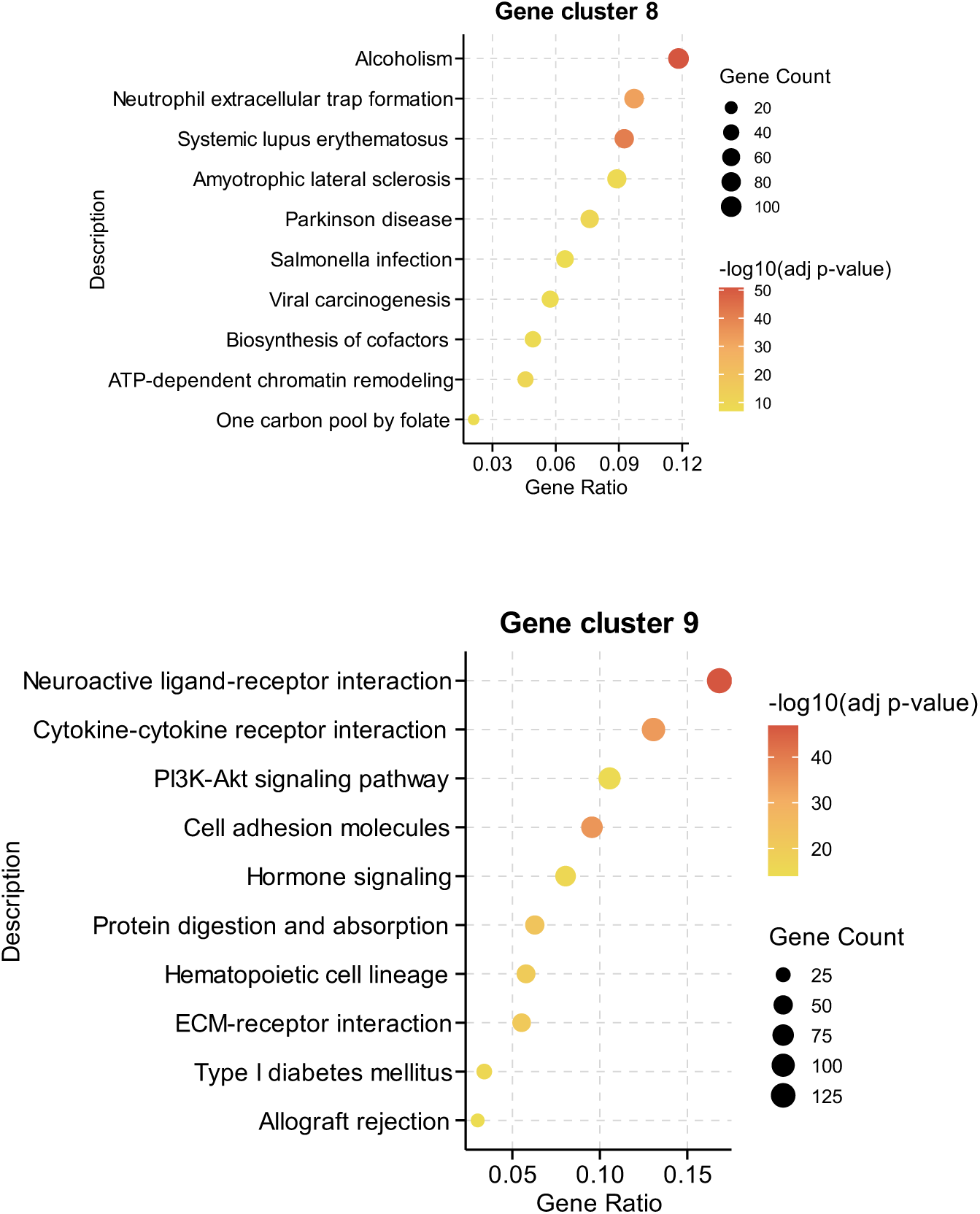
KEGG enrichment analysis of distinct gene clusters derived from gene embeddings generated by BulkFormer.

**Supplementary Figure 3.**
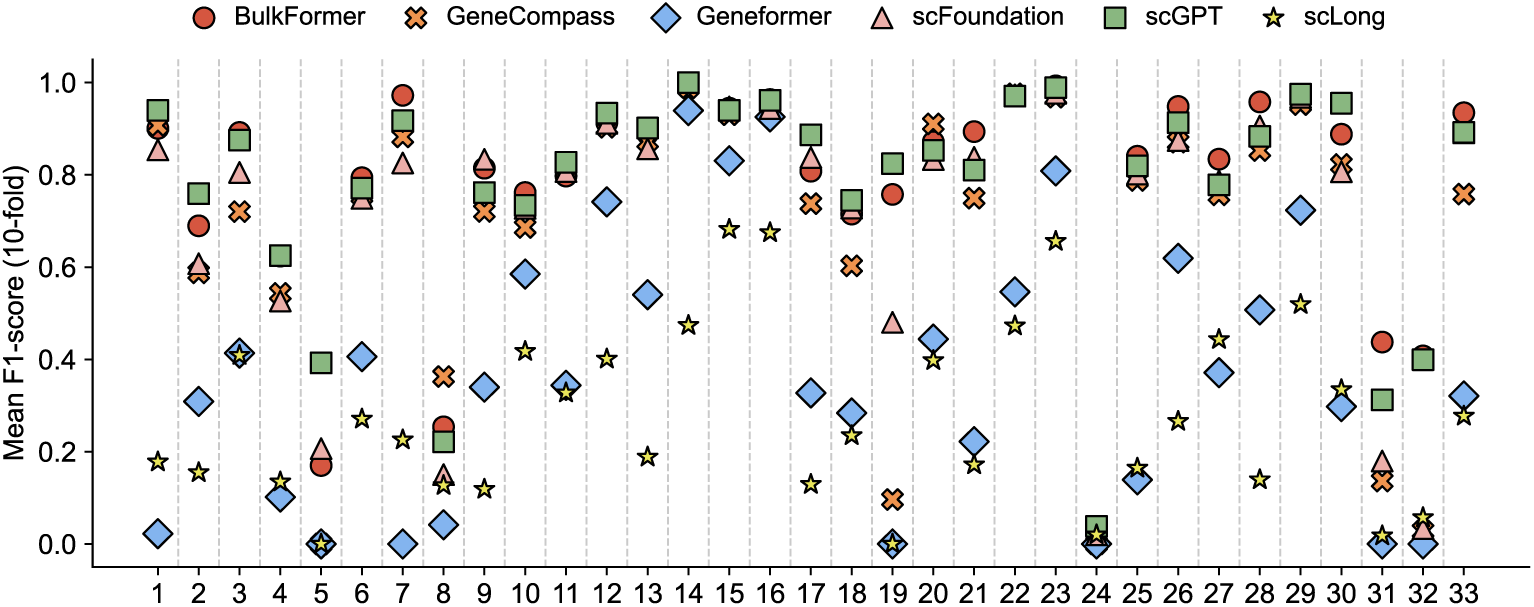
Classification performance of BulkFormer and baseline models on individual cancer subtypes.

**Supplementary Figure 4.**
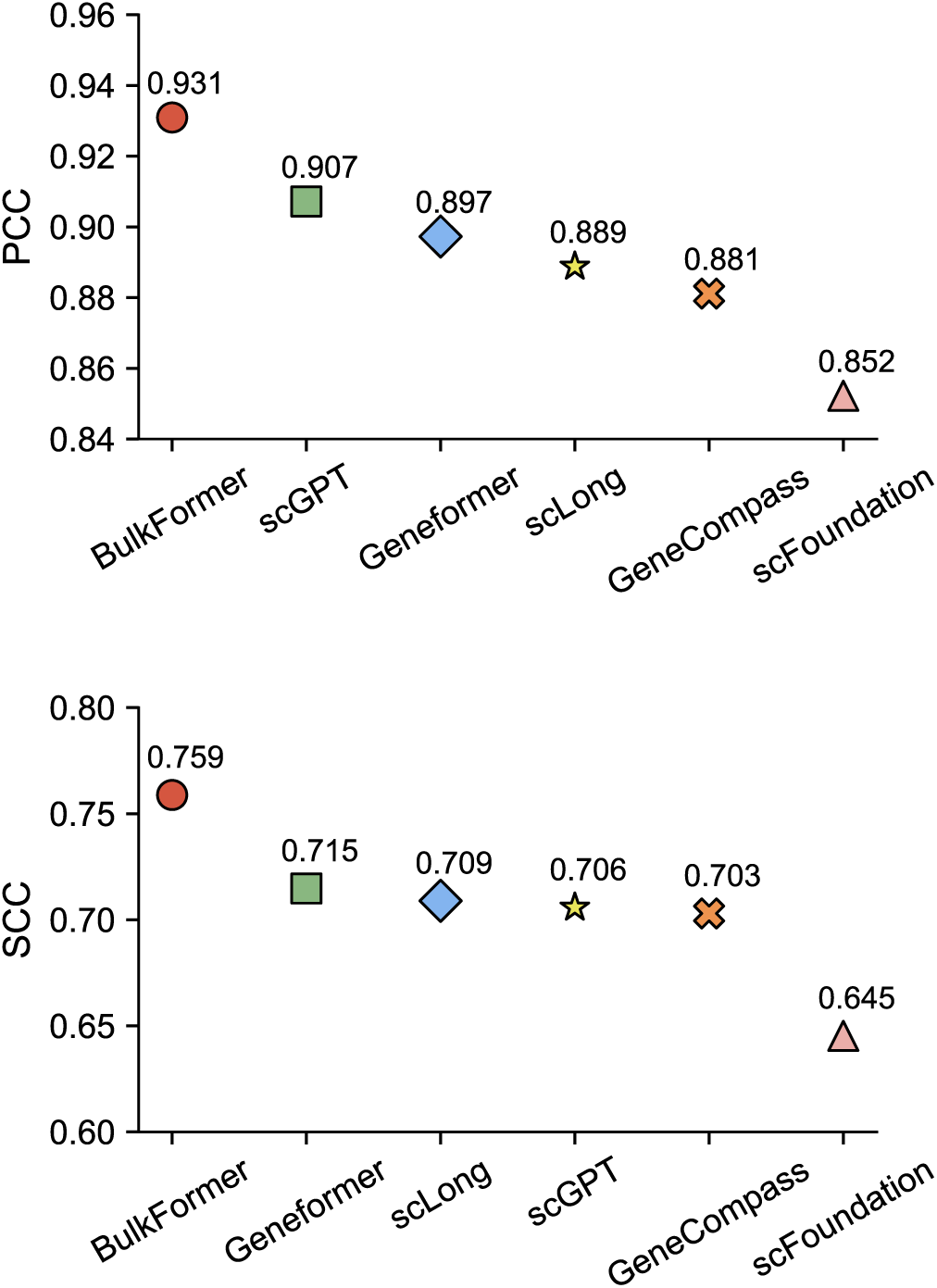
Performance comparison between BulkFormer and baseline models on the gene essentiality prediction task. PCC: Pearson correlation coefficient. SCC: Spearman correlation coefficient.

**Supplementary Figure 5.**
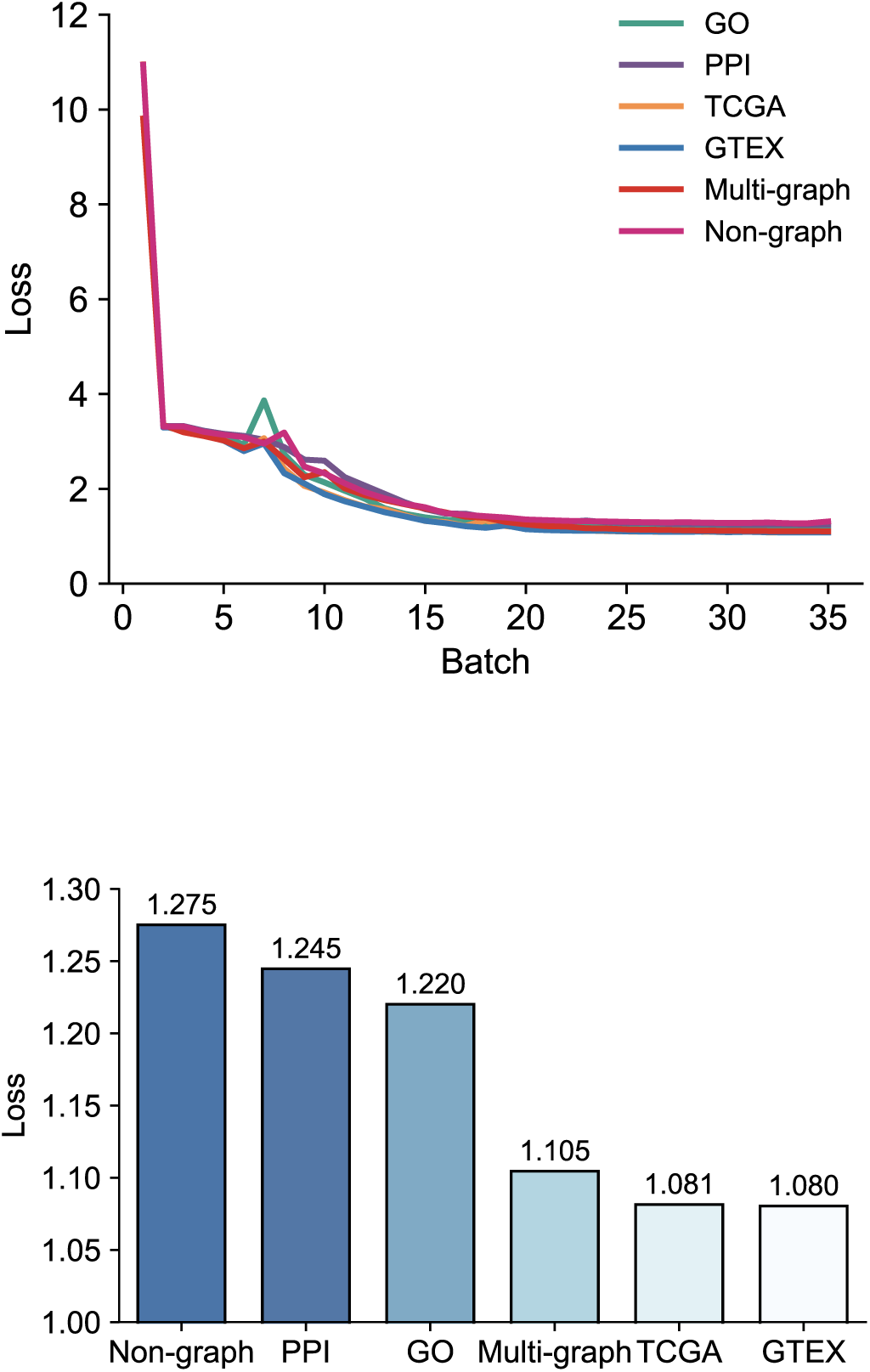
Comparison of BulkFormer performance using different biological knowledge graphs.

## Supplementary Tables

**Supplementary Table 1.**
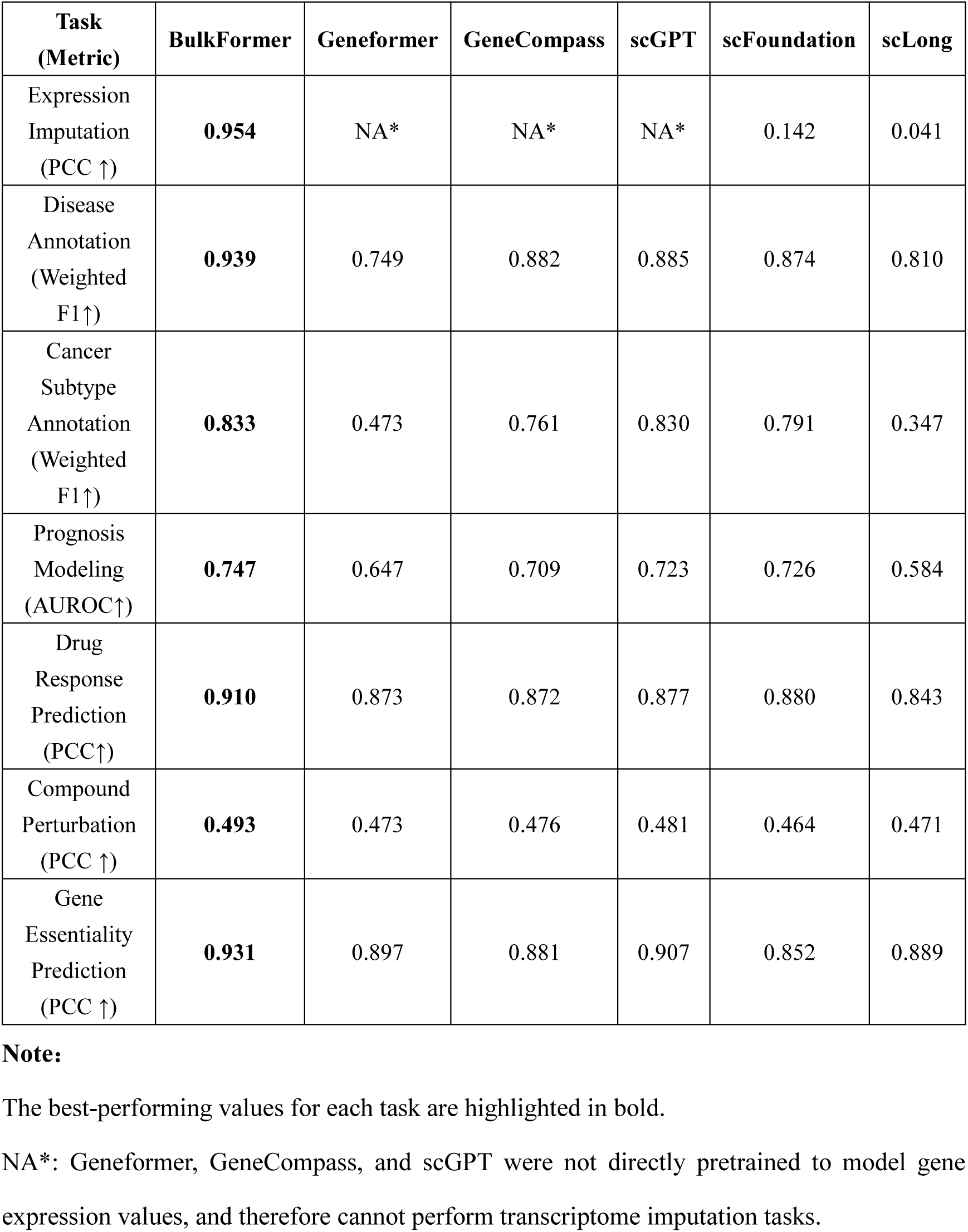
Performance comparison of BulkFormer and baseline models across six downstream tasks.

**Supplementary Table 2.** Detailed Comparison Between BulkFormer and scRNA-seq foundation models.

|  | **BulkFormer** | **Geneformer** | **scGPT** | **scFoundation** | **GeneCompass** | **scLong** |
| --- | --- | --- | --- | --- | --- | --- |
| Data size | ~0.5M | ~30M | ~10M | ~50M | ~126M | ~48M |
| Data type | Bulk | scRNA | scRNA | scRNA | scRNA | scRNA |
| Model size | ~150M | ~40M | ~50M | ~120M | ~132M | ~1,000M |
| Input value | Continuous value | Ranked value | Binned value | Continuous value | Continuous value and gene ID | Continuous value |
| Number of input genes (L) | 20,010 | 2,048 | ~2,000 | 19,264 | 2,048 | 27,874 |
| Embedding dim (D) | 640 | 512 | 512 | 768 | 768 | 200 |
| Training time* | ~96h | ~864h | NA* | NA* | ~2304h | ~16,128h |
| GPU device | NVIDIA A800 | NVIDIA V100 | NVIDIA A100 | NVIDIA Ampere GPU | NVIDIA A800 | NVIDIA A100 |

Note:
Training time\*: Here, training time refers to the average time required to complete one training epoch on a single GPU.
NA\*: The training times for scGPT and scFoundation were not reported in their publications.

**Supplementary Table 3.** Detailed information on 23 diseases collected from the DiSignAtlas database.

| ID | Name | Count |
| --- | --- | --- |
| 1 | COVID-19 | 2487 |
| 2 | Amyotrophic Lateral Sclerosis | 499 |
| 3 | Colorectal Carcinoma | 1499 |
| 4 | Type 1 Diabetes | 958 |
| 5 | Asthma | 3071 |
| 6 | Breast Cancer | 1871 |
| 7 | Psoriasis | 1003 |
| 8 | Alzheimer's Disease | 1328 |
| 9 | Parkinson's Disease | 1218 |
| 10 | Systemic Lupus Erythematosus | 2646 |
| 11 | Hepatocellular Carcinoma | 1750 |
| 12 | Chronic Obstructive Pulmonary Disease | 2041 |
| 13 | Sepsis | 657 |
| 14 | Influenza | 655 |
| 15 | Multiple Sclerosis | 1287 |
| 16 | Tuberculosis | 835 |
| 17 | Rheumatoid Arthritis | 1270 |
| 18 | Idiopathic Pulmonary Fibrosis | 1161 |
| 19 | Ulcerative Colitis | 3199 |
| 20 | Crohn's Disease | 2681 |
| 21 | Huntington's Disease | 764 |
| 22 | Acute Myeloid Leukemia (Aml-M2) | 671 |
| 23 | Schizophrenia | 1173 |

**Supplementary Table 4.** Detailed information on 33 cancer types collected from the TCGA database.

| ID | project | Name | count |
| --- | --- | --- | --- |
| 1 | ACC | Adrenocortical carcinoma | 77 |
| 2 | BLCA | Bladder Urothelial Carcinoma | 407 |
| 3 | BRCA | Breast Invasive Carcinoma | 1098 |
| 4 | CESC | Cervical Squamous Cell Carcinoma and Endocervical Adenocarcinoma | 306 |
| 5 | CHOL | Cholangiocarcinoma | 36 |
| 6 | COAD | Colon Adenocarcinoma | 288 |
| 7 | DLBC | Diffuse Large B-Cell Lymphoma | 47 |
| 8 | ESCA | Esophageal Carcinoma | 182 |
| 9 | GBM | Glioblastoma Multiforme | 165 |
| 10 | HNSC | Head and Neck Squamous Cell Carcinoma | 520 |
| 11 | KICH | Kidney Chromophobe | 66 |
| 12 | KIRC | Kidney Renal Clear Cell Carcinoma | 531 |
| 13 | KIRP | Kidney Renal Papillary Cell Carcinoma | 289 |
| 14 | LAML | Acute Myeloid Leukemia | 173 |
| 15 | LGG | Brain Lower Grade Glioma | 522 |
| 16 | LIHC | Liver Hepatocellular Carcinoma | 371 |
| 17 | LUAD | Lung Adenocarcinoma | 515 |
| 18 | LUSC | Lung Squamous Cell Carcinoma | 498 |
| 19 | MESO | Mesothelioma | 87 |
| 20 | OV | Ovarian Serous Cystadenocarcinoma | 427 |
| 21 | PAAD | Pancreatic Adenocarcinoma | 179 |
| 22 | PCPG | Pheochromocytoma and Paraganglioma | 182 |
| 23 | PRAD | Prostate Adenocarcinoma | 496 |
| 24 | READ | Rectum Adenocarcinoma | 92 |
| 25 | SARC | Sarcoma | 262 |
| 26 | SKCM | Skin Cutaneous Melanoma | 469 |
| 27 | STAD | Stomach Adenocarcinoma | 414 |
| 28 | TGCT | Testicular Germ Cell Tumors | 137 |
| 29 | THCA | Thyroid Carcinoma | 512 |
| 30 | THYM | Thymoma | 119 |
| 31 | UCEC | Uterine Corpus Endometrial Carcinoma | 181 |
| 32 | UCS | Uterine Carcinosarcoma | 57 |
| 33 | UVM | Uveal Melanoma | 79 |

**Supplementary Table 5.**
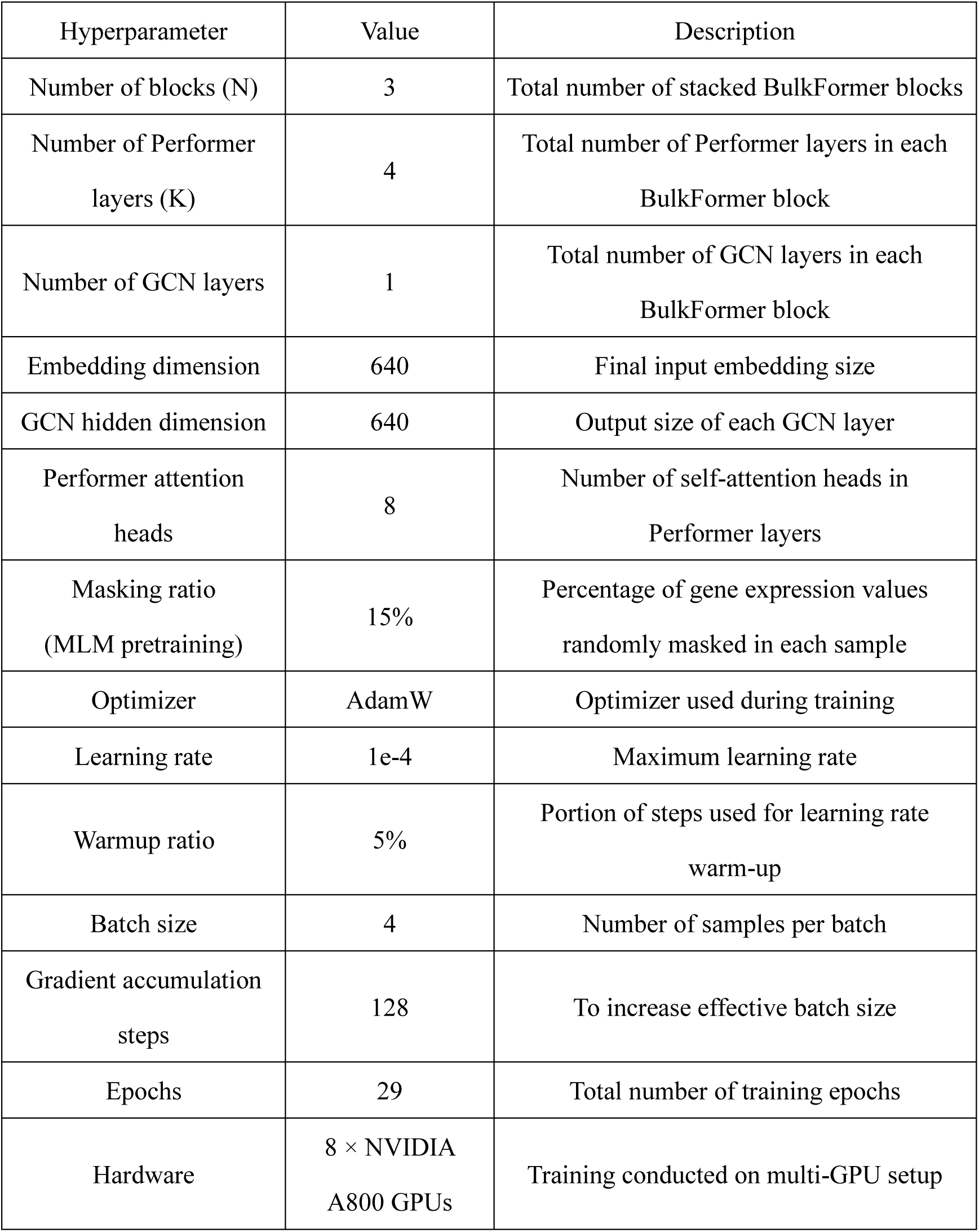
Hyperparameter settings used in BulkFormer model.

